# ROP INTERACTIVE PARTNER b interacts with RACB and supports fungal penetration into barley epidermal cells

**DOI:** 10.1101/750265

**Authors:** Christopher McCollum, Stefan Engelhardt, Lukas Weiss, Ralph Hückelhoven

**Author notes:** Author contributions: CM planned and performed experiments, prepared figures, interpreted results and wrote the manuscript. LW performed experiments. SE performed experiments, co-supervised CM, interpreted results and edited the manuscript. RH designed the study, planned experiments, supervised CM, interpreted results and wrote the manuscript. The author responsible for distribution of materials integral to the findings presented in this article in accordance with the policy described in the Instructions for Authors (www.plantphysiol.org) is: Ralph Hückelhoven. The project was funded in frame of research grants from the German Research Foundation to RH (DFG HU886/8 and SFB924).

## Abstract

RHO of Plants (ROP) G-proteins are key components of cell polarization processes in plant development. The barley (*Hordeum vulgare*) ROP protein RACB, is a susceptibility factor in the interaction of barley with the barley powdery mildew fungus *Blumeria graminis* f.sp. *hordei* (*Bgh*). RACB also drives polar cell development, and this function might be coopted during formation of fungal haustoria in barley epidermal cells. In order to understand RACB signaling during the interaction of barley with *Bgh*, we searched for potential downstream interactors of RACB. Here, we show that ROP INTERACTIVE PARTNER b (RIPb, synonym: INTERACTOR OF CONSTITUTIVE ACTIVE ROP b; ICRb) directly interacts with RACB in yeast and *in planta*. Over-expression of RIPb supports susceptibility of barley to *Bgh*. RIPb further interacts with itself at microtubules. However, the interaction with activated RACB takes place at the plasma membrane. Both, RIPb and RACB are recruited to the site of fungal attack around the neck of developing haustoria suggesting locally enhanced ROP activity. We further assigned different functions to different domains of the RIPb protein. The N-terminal coiled-coil CC1 domain is required for microtubule localization, while the C-terminal coiled-coil CC2 domain is sufficient to interact with RACB and to fulfill a function in susceptibility at the plasma membrane. Hence, RIPb appears to be localized at microtubules and is then recruited by activated RACB for a function at the plasma membrane during formation of the haustorial complex.

**One Sentence summary:** RIPb acts downstream of the powdery mildew susceptibility factor RACB of barley and influences susceptibility

## Introduction

The interaction of plants with powdery mildew fungi is a model for the biology of cell-autonomous responses to fungal parasites (Dörmann et al., 2014). The powdery mildew fungus *Blumeria graminis* f.sp. *hordei* (*Bgh*) is a biotrophic ascomycete specifically adapted to barley (*Hordeum vulgare*) and grows on the plant’s surface. In the beginning of its life cycle *Bgh* has to penetrate an epidermal cell in order to establish a haustorium for nutrient uptake (Hahn et al., 1997; Voegele et al., 2001) and to provide a surface for the translocation of virulence effector proteins into the host cell (Catanzariti et al., 2007). During all stages of fungal invasion, the epidermal host cell stays intact. Host cytosol and fungal haustorium are separated by the extrahaustorial matrix and the plant-derived extrahaustorial membrane (EHM).

Plant host cells polarize in very early phases of the interaction with fungi. A reorganization of the cytoskeleton was shown in different pathosystems, as well as the accumulation of peroxisomes, mitochondria, Golgi bodies and ER at the site of pathogen attack (Kobayashi et al., 1997; Takemoto et al., 2003; Koh et al., 2005; Takemoto et al., 2006; Fuchs et al., 2016). This is accompanied by relocation of the nucleus to the site of attack (Gross et al., 1993; Scheler et al., 2016). Polarization is considered important for effective defense, in particular for the focal formation of papilla or cell wall appositions, which requires localized deposition of callose, other cell wall glucans and phenolic compounds at the attempted penetration site (McLusky et al., 1999; Hückelhoven, 2007; Chowdhury et al., 2014). However, it is reasonable to assume, that host cell polarization is also important for successful pathogen establishment, for instance for the generation of the EHM (Scheler et al., 2016; Kwaaitaal et al., 2017).

ROP GTPases (RHO of Plants, also called RAC for rat sarcoma–related C3 botulinum toxin substrate) are small monomeric G-proteins that form a plant-specific RHO subfamily. ROPs can cycle between an actively signaling GTP-bound state and an inactive GDP-bound state and are crucial for polarity of diverse types of plant cells (Feiguelman et al., 2018). While activation is mediated by <u>G</u>uanosin Nucleotide <u>E</u>xchange <u>F</u>actors (GEF) enabling the exchange of GDP to GTP, inactivation is facilitated by <u>G</u>TPase <u>A</u>ctivating <u>P</u>roteins (GAP) which activate the intrinsic GTPase function of the G-protein, leading to GTP hydrolysis. ROPs seem to fulfill different functions depending on particular downstream factors called ROP-effectors. For instance *Arabidopsis thaliana* ROP2 suppresses light induced stomata opening by interacting with <u>R</u>OP <u>I</u>nteractive <u>C</u>RIB Motif Containing Protein7 (RIC7), which in turn interacts and inhibits the exocyst vesicle tethering complex subunit Exo70B1 (Hong et al., 2015). ROP2 is additionally involved in pavement cell lobe interdigitation by interacting with RIC4 for actin assembly in lobes and at the same time inhibiting RIC1 which organizes microtubules together with katanin KTN1 downstream of ROP6 (Fu et al., 2005; Lin et al., 2013). In these pathways, RIC proteins are considered scaffolds for connecting activated ROPs with downstream effector proteins in G-protein signaling.

Another class of downstream interactors are <u>R</u>OP <u>I</u>nteractive <u>P</u>artners (RIPs) first called <u>I</u>nteractors of <u>C</u>onstitutive Active <u>R</u>OPs (ICRs) (Lavy et al., 2007; Li et al., 2008). ICRs/RIPs represent a second class of plant-specific proteins connecting ROP signaling to downstream effectors. So far, little is known about these proteins. Arabidopsis knockout plants of RIP1/ICR1 have defects in pavement cell development, root hair development as well as root meristem maintenance showing an involvement of RIP1/ICR1 in different developmental processes. RIP1/ICR1 seems to be able to interact with different ROP proteins and was found to interact downstream with SEC3a of the exocyst complex and thereby possibly controlling the localization of the auxin transporter PIN1 (Lavy et al., 2007; Hazak et al., 2010; Hazak et al., 2014). Additionally it was reported, that RIP1 acts in pollen tube formation where it interacts with ROP1 at the plasma membrane of the pollen tube tip (Li et al., 2008). RIP3 (also called ICR5 or MIDD1 for <u>Mi</u>crotubule <u>D</u>epletion <u>D</u>omain1) plays a key role in xylem cell development in Arabidopsis. During the formation of the secondary cell wall in progenitor cells, RIP3 interacts with ROP11 and the kinesin KIN13A, which leads to local microtubule depletion and the formation of secondary wall pits (Mucha et al., 2010; Oda et al., 2010; Oda and Fukuda, 2012, 2013).

ROP GTPases also play a role as signaling components in plant defense (Ono et al., 2001; Chen et al., 2010). For instance, upon chitin perception, the receptor kinase CERK1 phosphorylates RacGEF1. RacGEF1 in turn activates RAC1, which supports immunity to *Magnaporthe oryzae* (Akamatsu et al., 2013).

The barley ROP protein RACB is involved in root hair outgrowth and controls asymmetric cell division of subsidiary cells in stomata development (Scheler et al., 2016). RACB and RACB-associated proteins influence arrays and stability of filamentous actin and the microtubule cytoskeleton (Opalski et al., 2005; Hoefle et al., 2011; Huesmann et al., 2012). Next to its function in polar cell development, RACB is also a susceptibility factor in the interaction with the powdery mildew fungus *Bgh*. Over-expression of constitutively activated RACB (CA RACB) enhances the penetration success of *Bgh* into barley epidermal cells, silencing of RACB leads to a decreased penetration rate (Schultheiss et al., 2002; Schultheiss et al., 2003; Hoefle et al., 2011). RACB’s function in susceptibility seems not to be dependent on defense suppression, but rather on the exploitation of developmental signaling mechanisms of the host (Scheler et al., 2016). A *Bgh* effector candidate, <u>ROP</u>-<u>I</u>nteractive <u>P</u>eptide1 (ROPIP1), binds directly to activated RACB. Expression of ROPIP1 in barley cells negatively influences microtubule stability and leads to an increased penetration rate of *Bgh* into barley epidermal cells (Nottensteiner et al., 2018). RACB further interacts with the class VI receptor-like cytoplasmic kinase <u>R</u>OP-<u>B</u>inding <u>K</u>inase1 (RBK1). Activated RACB supports *in vitro* kinase activity of RBK1, but RBK1 acts in resistance rather than susceptibility. This seems to be explained by the interaction of RBK1 with <u>S</u>-Phase <u>K</u>inase1-Associated (SKP1)-Like <u>P</u>rotein (SKP1-like), which is part of an E3-ubiquitin ligase complex and both RBK1 and SKP1-like can limit the abundance of the RACB protein (Huesmann et al., 2012; Reiner et al., 2015). Another interactor of RACB is the <u>M</u>icrotubule-<u>A</u>ssociated ROP <u>G</u>TPase <u>A</u>ctivating <u>P</u>rotein1 (MAGAP1), a CRIB-motif containing ROP-GAP. MAGAP1 and RACB recruit each other to the cell periphery and to the microtubule cytoskeleton, and MAGAP1 apparently counters the susceptibility effect of RACB, while silencing of MAGAP1 leads to increased susceptibility to *Bgh* (Hoefle et al., 2011).

In this study, we identified barley RIPb as another downstream interactor of RACB. We investigated the effect of RIPb on susceptibility by transient over-expression and RNAi knockdown of RIPb in single epidermal cells, and the interaction between RIPb and RACB by Yeast-Two-Hybrid assays and ratiometric bimolecular fluorescence complementation (BiFC). RIPb and RACB co-localize and presumably interact at the plasma membrane, at the microtubule cytoskeleton, and at the site of fungal invasion. To further investigate the structure-function relationship of RIPb, we tested a series of RIPb truncations regarding their function in the interaction of barley with *Bgh* and their role for protein-protein interaction.

## Results

### Identification of ICR/RIP proteins in barley

Previous studies have shown that ICR/RIP proteins are a class of proteins with little sequence similarity (Li et al., 2008). All ICR/RIP proteins identified so far in Arabidopsis contain an N-terminal QEEL motif and a C-terminal QWRKAA motif. These motifs are present in respective N- and C-terminal coiled-coil domains. Based on this, we performed bioinformatic analyses and identified three high confidence genes coding for ICR/RIP proteins in barley (Supplemental Fig. S1). It appears that in several grasses the first glutamic acid in the QEEL motif is exchanged to aspartic acid (QDEL). Because we did not observe a clear orthology to individually numbered *Arabidopsis thaliana* ICR/RIP proteins and phylogenetic analysis was ambiguous as well, we named the barley proteins RIPa/ICRa (HORVU3Hr1G087430), RIPb/ICRb (HORVU1Hr1G012460) and RIPc/ICRc (HORVU3Hr1G072880) (Supplemental Fig. S2). We also identified three ICR/RIP proteins in rice containing the QDEL motif as well as the QWRKAA motif (Os01g61760, Os05g03120 and OsJ_03509 (Yu et al., 2005)). Alignments of the barley ICRs/RIPs with the ICR/RIP proteins from rice and the five ICR/RIP proteins previously identified in Arabidopsis (ICR1/RIP1 (At1g17140), ICR2/RIP2 (At2g37080), ICR3/RIP5 (At5g60210), ICR4/RIP4 (At1g78430) and ICR5/RIP3/MIDD1 (At3g53350)) show little overall amino acid sequence conservation between the grasses and Arabidopsis, except for the conserved QD/EEL motif at the more N-terminal part and the QWRKAA motif at the more C-terminal part of the protein. The latter was shown to be necessary for ROP interaction (Lavy et al., 2007). The alignment also shows conservation of several lysine residues at the very C-terminus, which were shown before to be important for membrane localization of other ICR/RIP proteins (Li et al., 2008) (Supplemental Fig. S1).

Phylogenetic analysis shows that HvICRa/HvRIPa and HvICRb/HvRIPb are more closely related to each other, than to HvICRc/ HvRIPc, which is located on an independent branch of the tree (Supplemental Fig. S2). Each one ICR/RIP from rice (*Oryza sativa* ssp. *japonica*) and *Brachypodium distachyon* appear to be orthologous to HvICRa/HvRIPa, HvICRb/HvRIPb and HvICRc/HvRIPc respectively. (Supplemental Fig. S2).

### RIPb influences susceptibility of barley to *Bgh*

Semiquantitative reverse transcription PCR showed that all three barley ICRs/RIPs are transcribed in whole leaves and the epidermis, with *RIPb* showing the highest RNA levels and *RIPa* being only weakly expressed. Samples from inoculated leaves showed that *Bgh* infection does not alter expression of any of the three barley *ICRs/RIPs* (Supplemental Fig. S3). To investigate, if one of the ICRs/RIPs influences susceptibility of barley to *Bgh*, we tested the penetration efficiency of *Bgh* into transiently transformed epidermal cells. We introduced either an over-expressing construct under control of the CaMV35S promotor or a posttranscriptional gene-silencing construct into these cells. Over-expression of RIPa or RIPc had no significant effect on susceptibility (Supplemental Fig. S4A) (Barley ICR/RIP proteins are called RIPs from now on for reasons of simplicity). Over-expression of RIPb however, significantly and consistently increased the penetration rate of *Bgh* into transformed cells by 22%, compared to cells transformed with the empty vector control (Fig. 1A). RNA interference (RNAi)-mediated silencing of RIPb, did not significantly change the penetration rate of *Bgh* into the transformed cells (Fig. 1B).

**Figure 1.**
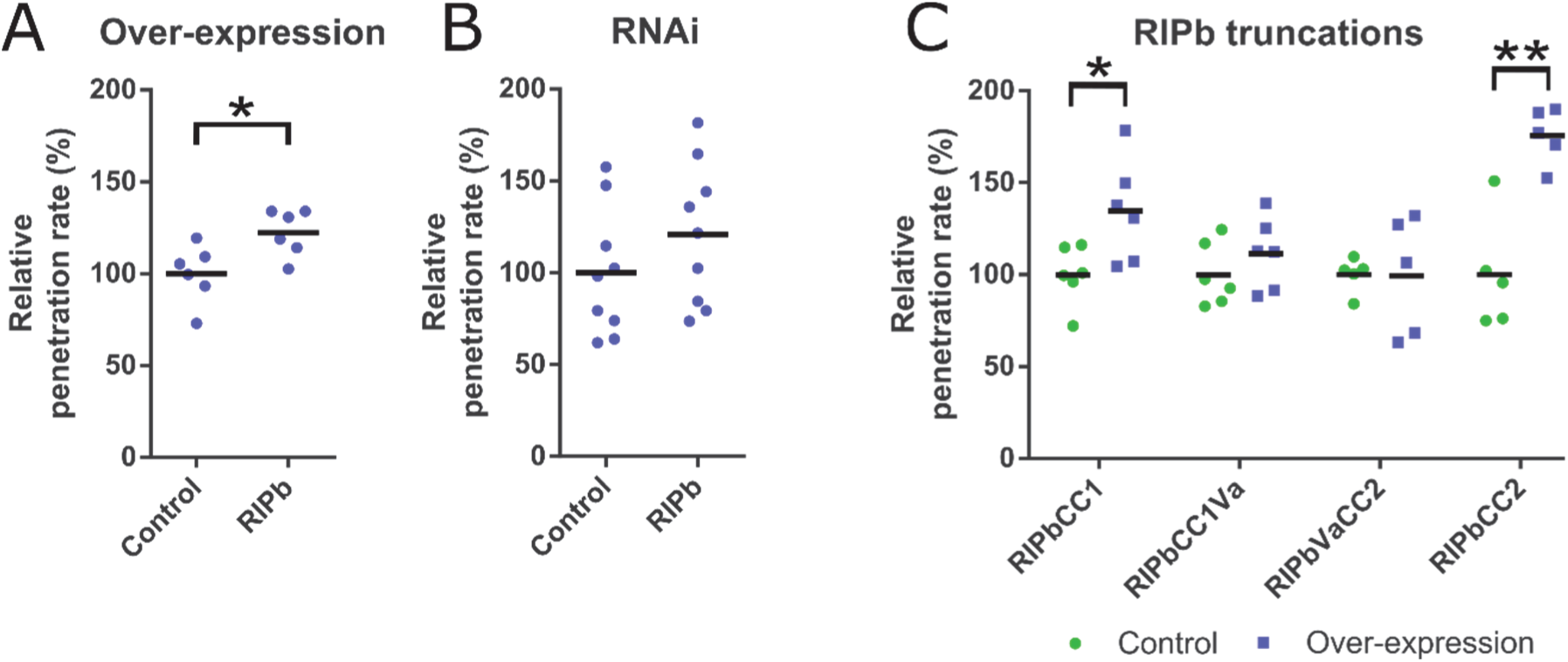
Effect of RIPb on the interaction of barley and *Bgh* was tested by biolistic transformation of epidermal cells of 7 days old barley leaves and determining the penetration rate of *Bgh* into the transformed cells 24 h after inoculation. Over-expression constructs for *RIPb* (A) as well as an RNAi silencing construct for *RIPb* (B) and over-expression constructs for *RIPb* truncations were introduced (C). As control, the respective empty vectors were used. Values represent the mean values of results of individual experiments (n≥5) relative to the mean of the respective control set as 100 %. One asterisk indicates significance *P* < 0.05; two asterisk indicate significance *P < 0*.*01*, Students t-test.

### RIPb interacts with RACB

In order to determine the subcellular localization of RIPb, we transiently expressed an YFP-tagged fusion protein of RIPb in single epidermal cells via biolistic transformation. YFP-RIPb was detected in the cytoplasm, at cytoskeleton structures and at the cell periphery. Co-expression experiments showed partial co-localization of YFP-RIPb and the barley microtubule marker MAGAP1-Cter at cortical microtubules (Fig. 2). The RFP-MAGAP1-Cter marker does not interact with ROPs because it lacks the ROP-interacting CRIB and the GAP domains but still contains the microtubule interacting <u>C</u>-<u>ter</u>minal part of MAGAP1 (Hoefle et al., 2011). Recorded images further confirmed that YFP-RIPb is also present in the cytosol, where it co-localized with additionally co-expressed soluble CFP, and at the cell periphery or plasma membrane (Fig. 2). Co-expression with constitutively activated RACB-G15V (CA RACB) resulted in depleted cytosolic localization of YFP-RIPb, when compared to soluble red fluorescing mCherry. At the same time, microtubule localization and cell-peripheral localization was still clearly detectable. Cytoplasmic depletion of YFP-RIPb was not observed when we co-expressed dominant negative RACB-T20N (DN RACB) (Fig. 3A). This change in RIPb localization might be best explained if RACB recruits RIPb to the cell periphery/plasma membrane. To test this, we co-expressed YFP-RIPb with the plasma membrane marker pm-rk (Nelson et al., 2007; Weis et al., 2013), either alone or in presence of CA RACB (Supplemental Fig. S5). YFP-RIPb alone showed some overlapping signal with pm-rk, but the peak in the signal profile was slightly displaced due to additional cytosolic signal (Supplemental Fig. S5A). However, in presence of CA RACB we recorded a shift of the YFP-RIPb peak towards the peak of the plasma membrane marker (Supplemental Fig. S5B).

**Figure 2.**
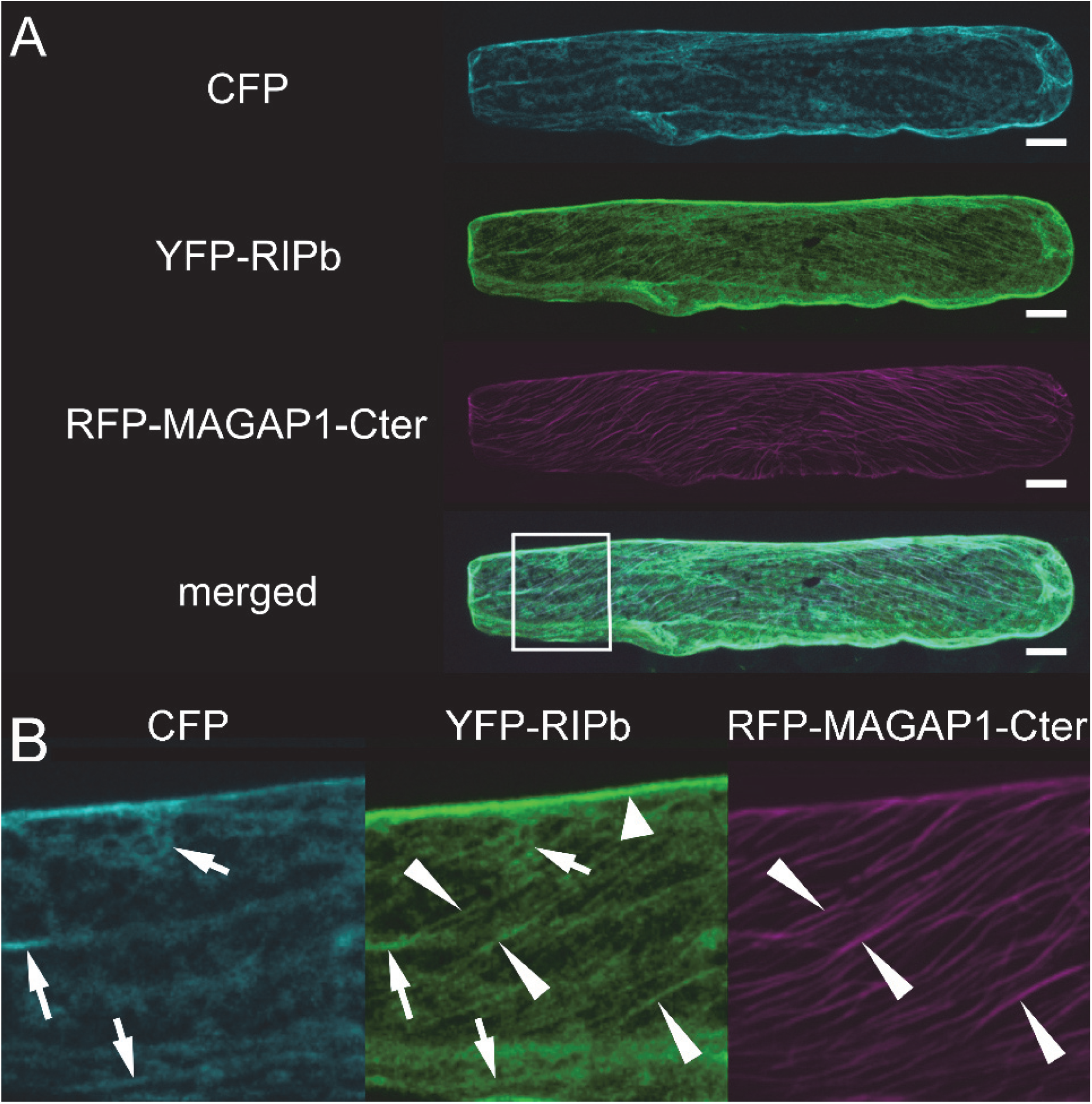
Subcellular localization of RIPb *in planta*. (A) Barley epidermal cell was transiently co-transformed with CFP as a cytosolic marker, YFP-tagged RIPb (YFP-RIPb) and RFP-MAGAP1-Cter as a microtubule marker. Image shows z-stacks of optical sections at the confocal laser scanning microscope of upper half of the cell. Bars represent 20µm. (B) Magnification of the part of the cell highlighted in A (merged channel). Please note the YFP-RIPb signal in the cytoplasm (arrows, compare to CFP), at the upper cell periphery (big arrowhead, highlighted only in the YFP-RIPb channel) and at microtubules (long narrow arrowheads, compare to RFP-MAGAP1-Cter). Brightness of the images was equally increased for displaying purposes. Pictures represent z-stacks of 10 confocal sections of each 2 µm increment.

**Figure 3.**
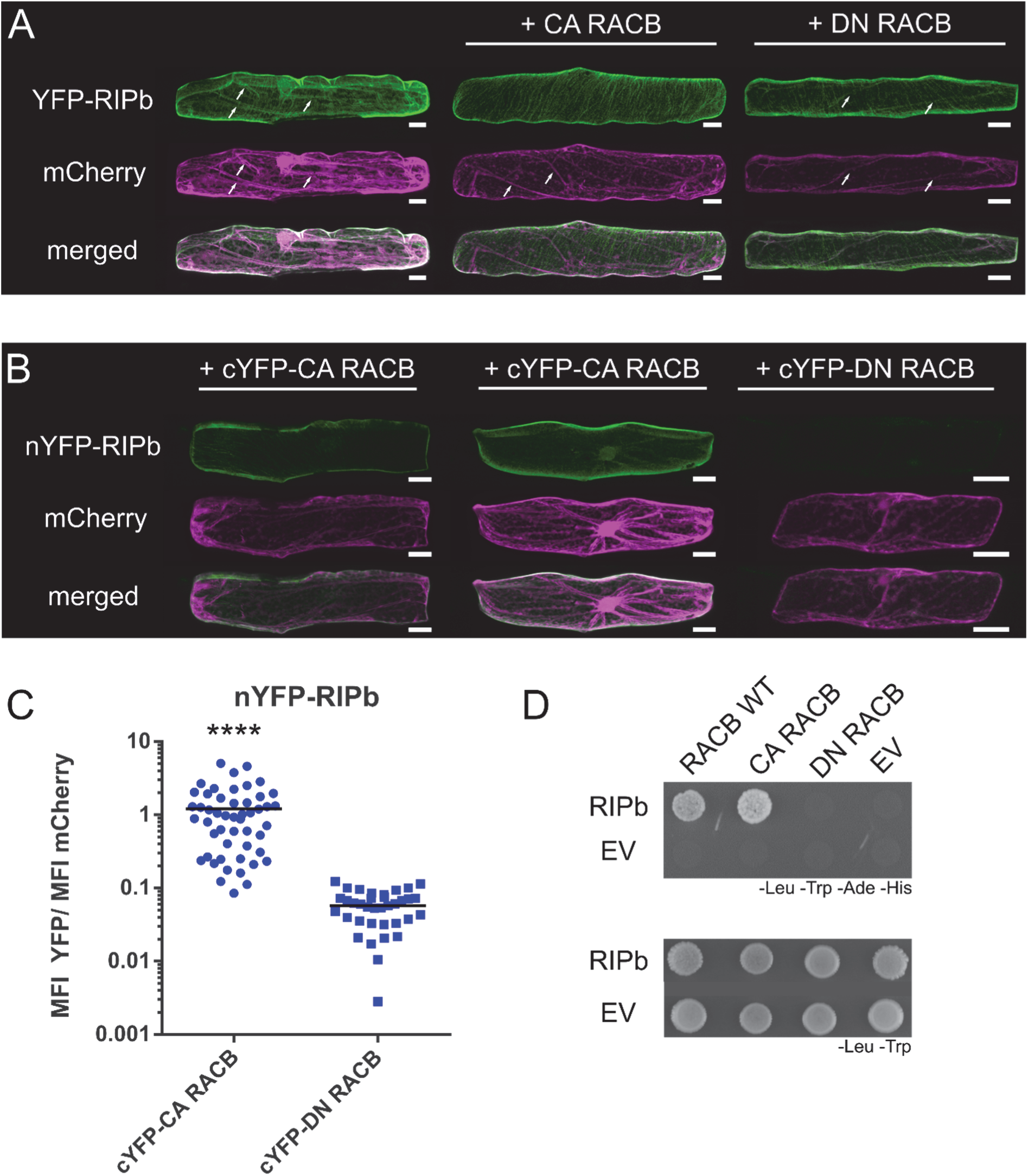
RACB and RIPb interact in yeast and in planta. (A) Single epidermal cells were transiently transformed by particle bombardment. YFP-RIPb and cytosolic transformation marker mCherry were expressed alone or co-expressed with constitutively activated RACB (CA RACB) or dominant negative RACB (DN RACB), respectively. Images were taken 24 hours after bombardment (hab) and show representative z-stacks of XY optical sections of the upper half of the cells. White arrows show cytosolic strands. White bars correspond to 20µm. (B) For BiFC experiments, protein fusions of RIPb, CA RACB and DN RACB with split-YFP tags were coexpressed. (B) images were taken 24 hab. Images show z-stacks of 10 optical sections of the upper half of the cells. White bars correspond to 20µm. (C) For quantification of BiFC experiments images were taken with constant settings and signal intensity (Mean Fluorescence Intensity, MFI) was measured over a region of interest at the cell periphery. The ratio between YFP and mCherry fluorescence signal was calculated. The figure shows one out of two replicates with similar results. For each replicate >30 cells were measured. (D) RIPb was tested in a Yeast-Two-Hybrid assay for its interaction with barley wild type RACB (RACB WT), CA RACB and DN RACB. As control the interaction with the respective empty vectors (EV) was tested. For identification of interactions SD medium lacking leucine (-Leu), tryptophan (-Trp), adenine (-Ade) and histidine (-His) was used. For identification of transformed cells SD medium lacking leucine and tryptophan was used.

This supports that RIPb gets recruited to the plasma membrane by activated RACB, but also shows that RIPb itself localized close to or attached to the plasma membrane. Ratiometric Bimolecular Fluorescence Complementation (BiFC) experiments further supported the interaction of RIPb with RACB. YFP fluorescence was reconstituted when nYFP-RIPb and cYFP-CA RACB were co-expressed in leaf epidermal cells (Fig. 3B, C). By contrast, co-expression of nYFP-RIPb and cYFP-DN RACB did not result in clear BiFC and the strength in signals were on average less than 10% of the signals recorded for the interaction with CA RACB (Fig. 3B, C). We observed the complemented CA RACB-RIPb YFP complex signals either exclusively at the plasma membrane or at cortical microtubules and the plasma membrane but hardly in the cytosol, when compared to mCherry (Fig. 3B). We further confirmed a direct interaction between both wild type RACB (RACB WT) and CA RACB with RIPb (Fig. 3D), respectively, in yeast. These experiments together suggest a direct interaction between RIPb and RACB *in planta*.

### RIPb truncations show distinct subcellular localization and function

All predicted ICR/RIP proteins from *H. vulgare, O. sativa, A. thaliana* and *B. distachyon* contain an N-terminal coiled-coil-(CC) domain with the QD/EEL motif as well as a C-terminal CC-domain with the QWRKAA motif (Supplemental Fig. S1; Fig. 4A). Based on this and with regard to previous studies (Mucha et al., 2010), we created RIPb truncations based on the first CC-domain (CC1), the central variable region (Va) and the second CC-domain (CC2). In yeast, only constructs containing the CC2 domain and hence the QWRKAA motif interacted with CA RACB as it was shown before for the interaction of Arabidopsis ROPs and ICRs/RIPs (Fig. 4B) (Lavy et al., 2007; Mucha et al., 2010). RIPbCC2 was also able to interact with CA RACB but not with DN RACB in BiFC assays and this interaction took place at the plasma membrane (Supplemental Fig. S7). To confirm this, we again co-expressed a YFP fusion of RIPbCC2 with the plasma membrane marker pm-rk either alone or in presence of CA RACB (Supplemental Fig. S5). Similar to full length RIPb, YFP-RIPbCC2 showed co-localization with pm-rk but the peak was slightly shifted in the signal profile probably due to additional cytosolic signal (Supplemental Fig. S5C). Co-expression with CA RACB again shifted the signal towards the plasma membrane, resulting in an overlay of the two peaks in the signal profile (Supplemental Fig. S5D). RIPb was also able to interact with itself in yeast. The Va-region may be important for this, since only full length RIPb and truncations containing this region were able to interact in yeast (Fig. 4C). In order to look for specific subcellular localizations *in planta*, we created YFP-tagged protein fusions of these truncations. Like YFP-RIPbCC2, YFP-RIPbVaCC2 localized strongly to the cell periphery, presumably the plasma membrane, with weak cytosolic background (Fig. 4D). YFP-RIPbCC1Va was located in the cytosol and at the microtubules. Additionally, co-expression experiments with YFP-RIPbCC1Va and CA RACB show that YFP-RIPbCC1Va was not positioned at the cell periphery despite the presence of CA RACB (Supplemental Fig. S6), suggesting that the CC2 domain is necessary for the recruitment by CA RACB. However, YFP-RIPbCC1 and YFP-RIPbVa were exclusively detected in the cytosol (Fig. 4D). Hence, both the CC1 domain and the Va domain appear to be required but not sufficient for microtubule association. Double mutation of D85 and E86 of the QDEL motif did not lead to a loss of microtubule localization (Supplemental Fig. S6B). The QDEL motif itself might therefore not be necessary for microtubule localization. Since the Va domain was required for dimerization and microtubule association, RIPb might localize to the microtubules as a dimer or oligomer. This was further supported because BiFC-signals recorded after co-expression of nYFP-RIPb with cYFP-RIPb occur exclusively at the microtubules and show less cytosolic background (Fig. 5A), when compared to YFP-RIPb alone, which may be detected both in its monomeric and its dimeric/oligomeric form (Fig. 2). Quantification of reconstituted YFP fluorescence showed significantly stronger signal intensities for the nYFP-RIPb/cYFP-RIPb homodimer compared to the co-expression of MAGAP1-nYFP with cYFP-RIPb, which are not supposed to interact and used as a negative control. As a positive control for the negative control, we co-expressed MAGAP1-nYFP with cYFP-CA RACB, which again showed high YFP complementation signals (Fig. 5A, B). Signal quantification showed high signal overlap between the complemented YFP fluorescence signal and microtubule marker RFP-MAGAP1-Cter over a linear region of interest (Fig. 5C, D). Since there appeared to be little cytosolic background in the nYFP-RIPb/cYFP-RIPb BiFC images, we measured the ratio between microtubule and cytosolic signal within each cell. We then compared signal ratios within YFP-RIPb expressing cells and those expressing the nYFP-RIPb/cYFP-RIPb BiFC pair. As a positive control for microtubule localization, we also measured the signal ratio for the microtubule marker RFP-MAGAP1-Cter. The data show that the microtubule/cytosol ratio for nYFP-RIPb/cYFP-RIPb BiFC was far higher than that of YFP-RIPb, indicating a more exclusive microtubule localization of the di/oligomer (Fig. 5E). In fact, the signal ratio for nYFP-RIPb/cYFP-RIPb was similar to that of the microtubule marker RFP-MAGAP1-Cter.

**Figure 4.**
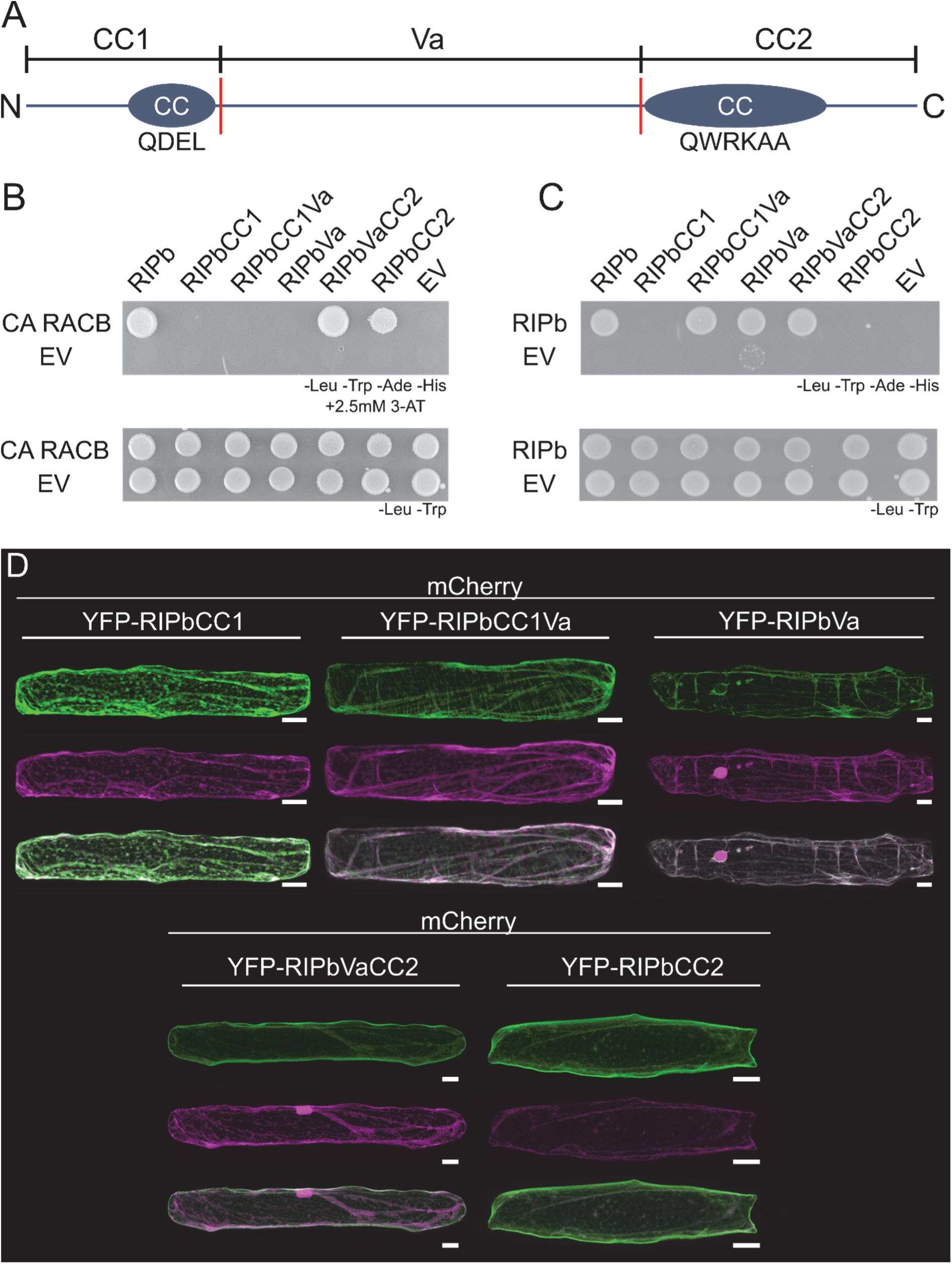
Structure function relationship of RIPb. (A) Domain structure and truncations of RIPb. The CC1-domain stretches from amino acid (aa) 1 to 132 and contains the N-terminal coiled-coil domain with the QDEL motif (CC, circles). The variable region (Va) starts at aa 133 at ends at aa 420. The CC2-domain stretches from aa 421 to the end at aa 612. The CC2-domain also represents a coiled-coil structure and contains the QWRKAA motif. (B) RIPb truncations were tested in Yeast-Two-Hybrid assays for their interaction with constitutively activated RACB (CA RACB) or RIPb (shown in C), respectively. As controls, the interaction with the respective empty vector (EV) was tested. For identification of interactions SD medium without leucine (-Leu), tryptophan (-Trp), adenine (-Ade) and histidine (-His) was used, together with 2,5mM 3-amino triazol to reduce background growth in the combinations containing the RIPbVa truncation. For identification of transformed cells SD medium without leucine and tryptophan was used. (D) Single epidermal cells were transiently transformed with different RIPb truncations tagged to YFP. Images show z-stacks of 15 optical sections of upper half of cells. White bars correspond to 20µm.

**Figure 5.**
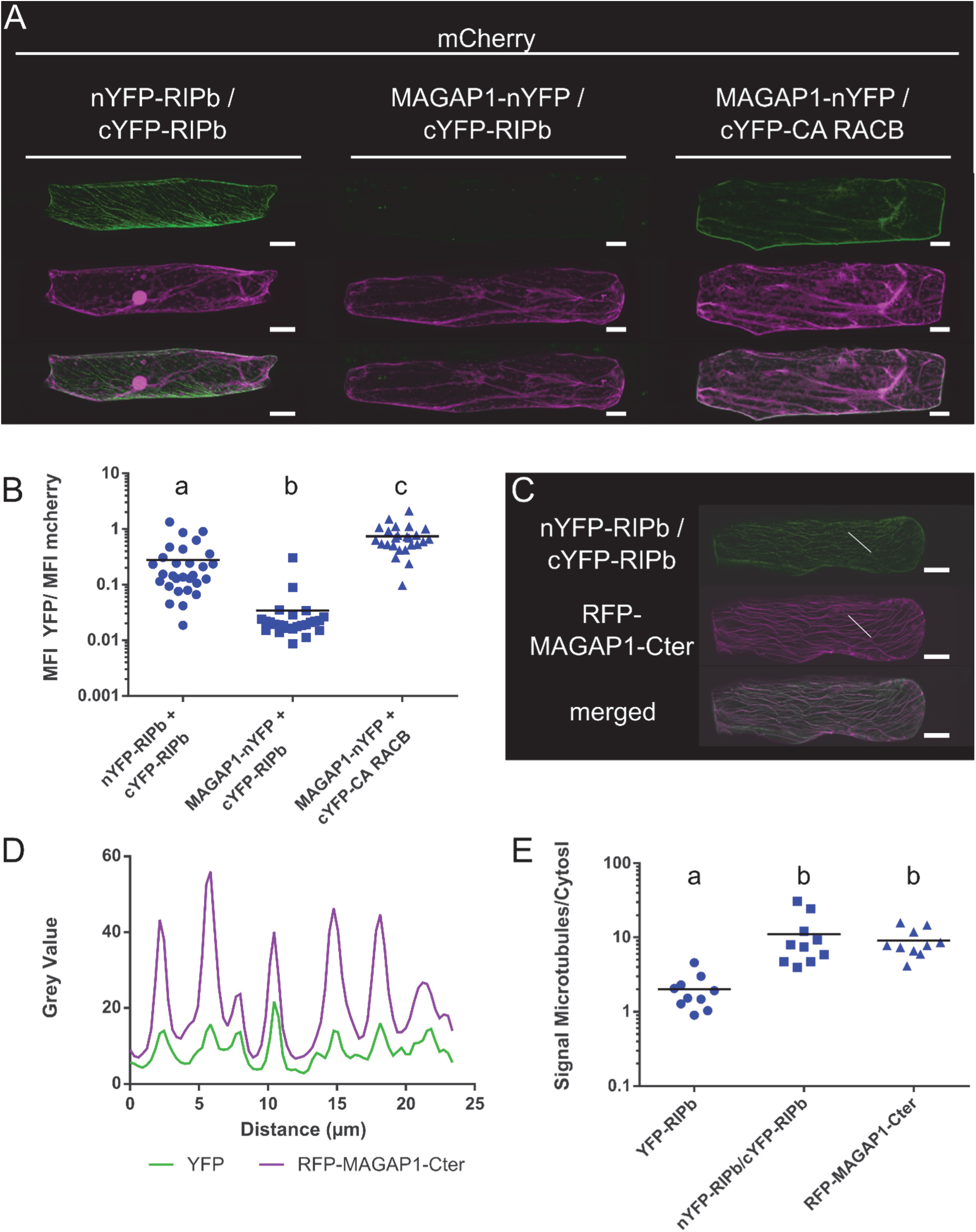
RIPb-RIPb interaction at microtubules. (A) Single epidermal cells were transiently transformed by particle bombardment with split-YFP constructs in the combination nYFP-RIPb and cYFP-RIPb, MAGAP1-nYFP and cYFP-RIPb as well as MAGAP1-nYFP and cYFP-RIPb. (B) Quantification of BiFC signals from images were taken with constant settings. Signal intensity (Mean Fluorescence Intensity, MFI) was measured over a region of interest at the cell periphery. The ratio between Split-YFP and free mCherry signal was calculated. Letters indicate significance by one-way ANOVA (Tukey’s multiple comparison test). Each signals were measured in 10 cells for each construct. Letters indicate significance by one-way ANOVA (Tukey’s multiple comparison test). (C) Co-expression of nYFP-RIPb and cYFP-RIPb with RFP-MAGAP1-Cter. Image brightness was equally increased for displaying purposes, but signal intensities over a region of interest (white line) were measured using original data (see D). White bars correspond to 20µm. (E) Ratio between microtubule signal and cytosolic signal was measured for YFP-RIPb alone, BiFC signal for RIPb-RIPb interaction and the microtubule marker RFP-MAGAP1-Cter. Microtubule and cytosolic signals were measured at three different regions of interest in the cell. Pictures represent z-stacks of 10 confocal sections of each 2 µm increment. Measurements were made in single optical section of the z-stack. Note the logarithmic scales.

Results from Lavy et al. (2007) and Mucha et al. (2010) suggest, that ICRs/RIPs lacking a functional QWRKAA motif lose the ability to interact with ROPs and that either CC1 or CC2 domains can bind to further downstream signaling components. This indicates that RIPb might be able to fulfill a ROP signaling function through one of these domains. To test the functionality of RIPb truncations, we tested their effect on penetration success of *Bgh* on barley. Interestingly over-expression of RIPbCC2 strongly increased susceptibility by about 75% (Fig. 1C). In contrast, over-expression of the CC2-domain of RIPa did not lead to a significant increase in susceptibility (Supplemental Fig. S4B). The effect of RIPbCC2 completely disappeared when we expressed RIPbVaCC2, containing additionally the Va-domain. The CC1-domain alone also increased susceptibility by about 35% and this effect was also reduced when we expressed the longer RIPbCC1Va truncation (Fig. 1C). In order to investigate the possible influence of protein levels on the influence on susceptibility, we measured the fluorescence intensity of YFP-tagged fusion proteins relative to an internal mCherry control in single transformed cells. We found that fluorescence intensities of full length RIPb and YFP-RIPbVaCC2 were lower than that of RIPbCC1 and RIPbCC2 (Supplemental Fig. S8). Hence, protein expression levels might have influenced the strength of induced susceptibility but CC1 and particularly CC2 alone were sufficient to support fungal penetration success.

Oda et al. (2010) showed that in Arabidopsis RIP3/ICR5/MIDD1 is involved in local microtubule depolymerization during xylem cell development. Microtubule depletion might also influence the outcome of the interaction of barley and *Bgh*. Indeed, we recently showed that barley RIPa influences microtubule organization when co-expressed with barley RAC1 and MAGAP1 (Hoefle et al., 2020). To see if this could be the case for RIPb in barley, we co-expressed RIPb and RIPbCC2, respectively, with the microtubule marker RFP-MAGAP1-Cter and evaluated microtubule organization as described by Nottensteiner et al. (2018). In the empty vector control, we found 68% of microtubules in a well-organized parallel state, while about 17% of cells show disordered but intact microtubules and the rest of the cells showed fragmented microtubules (Supplemental Fig. S9). In cells expressing RIPb or RIPbCC2 we observed a similar pattern. The amount of fragmented microtubules was a little lower in cells expressing RIPb and a little higher in cells expressing RIPbCC2, but no statistically significant changes were observed.

### RACB and RIPb co-localize at the site of fungal attack

Since RIPb and RACB can interact *in planta* and both proteins can influence barley susceptibility to *Bgh*, we wanted to know whether RIPb and RACB would localize to the sites of fungal penetration. Therefore, we transiently co-expressed YFP-RIPb and CFP-RACB in single epidermal cells and inoculated the leaves with conidia of *Bgh*. At 24 h after inoculation, we observed ring-like accumulations of both YFP-RIPb and CFP-RACB at the site of fungal penetration around the haustorial neck. Cytosolic mCherry appeared also in regions at the site of attack but was less spatially confined than YFP-RIPb and CFP-RACB (Fig. 6A). We observed even more pronounced fluorescence at infection sites, when YFP-tagged RIPb was co-expressed with CFP-CA RACB. In this context, we detected clear accumulation of RIPb and CA RACB at the site of fungal penetration, though independent of the outcome of the penetration attempt. If the penetration was successful, a clear ring-like localization pattern around the haustorial neck could be observed. However, if the fungal penetration was not successful we detected a more fringed accumulation of both proteins, possibly representing membrane domains around papilla protrusions (Fig. 6B). Since RIPbCC2 had a stronger influence on fungal penetration success than full length RIPb, we also imaged YFP-RIPbCC2 when co-expressed with CFP-CA RACB. Interestingly, there was a very strong localization of both proteins around the haustorial neck region in penetrated cells, but also in some instances at sites of repelled fungal attempts (Fig. 6C). The ring-like accumulation of RIPbCC2 around the haustorial neck was also visible later at 48 hours after the inoculation (Fig. 6D). There was also constantly local aggregation of cytoplasm at the sites of attack, but measurements of the ring-like YFP-RIPbCC2 fluorescence showed signal intensities were clearly more confined to the cell periphery compared to cytosolic mCherry fluorescence (Fig. 6E).

**Figure 6.**
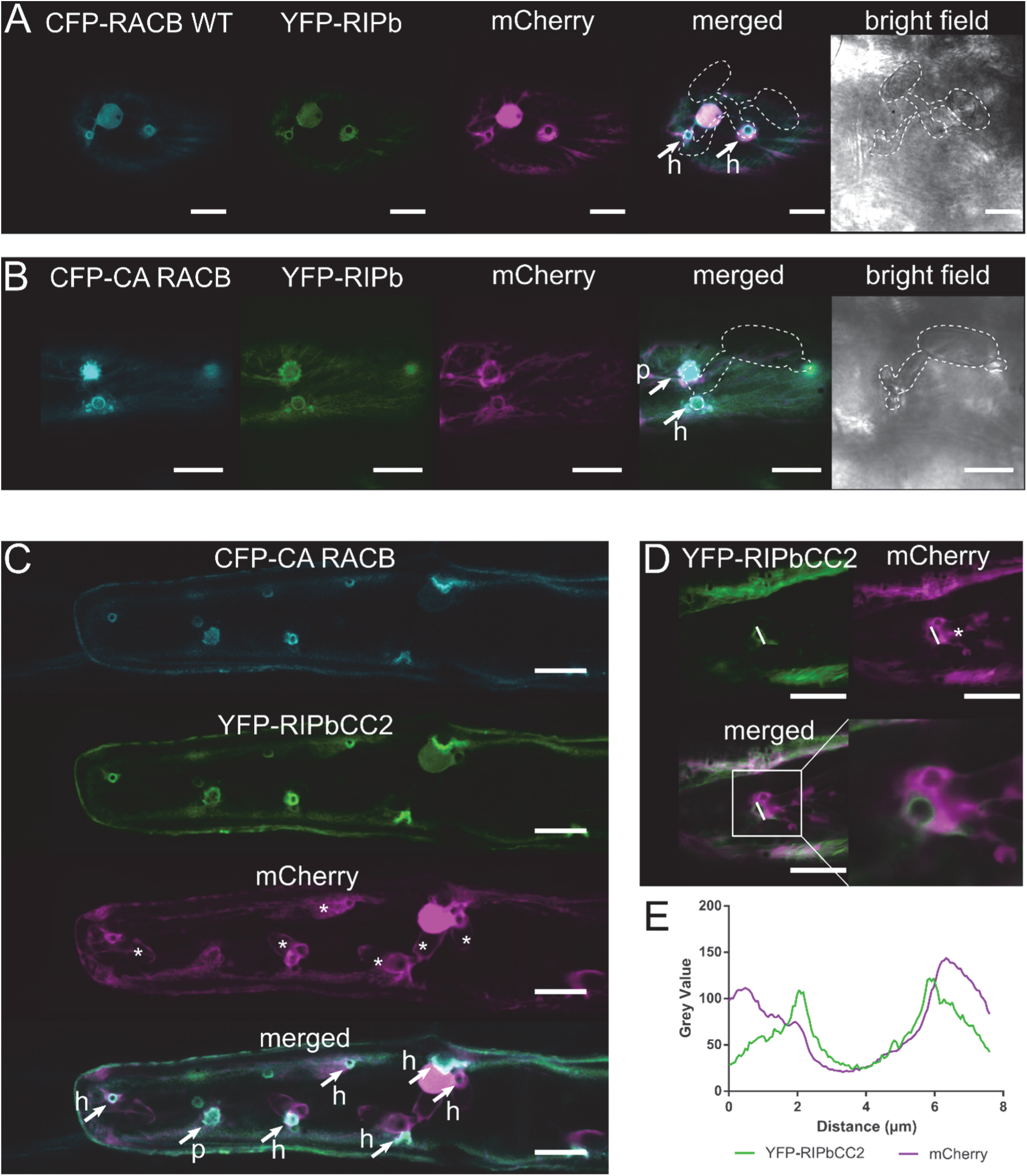
RIPb and RACB co-localize at sites of fungal attack. (A) Transiently transformed epidermis cells were inoculated with *Bgh*. YFP-RIPb co-localizes with CFP-RACB WT as well CFP-CA RACB (shown in B) at the site of fungal attack at 24 hours after inoculation. mCherry was used as a cytosolic marker. Confocal laser scanning recordings show z-stacks of the upper part the cells (5 confocal sections of each 2 µm increment). Transmission channel images show a single optical section. (C) YFP-RIPCC2 co-localizes with CFP-CA RACB 24 hours after inoculation at the site of fungal attack. mCherry was used as a cytosolic marker. Arrows mark sites of fungal penetration attempts that either succeeded with formation of a haustorium (h) or failed in a non-penetrated papilla (p). Asterisks indicate haustorial bodies. (D) Single epidermal cells were transiently transformed with YFP-RIPbCC2 and mCherry. Images were taken 48 hours after inoculation with *Bgh*. Signal intensities at the haustorial neck over the region of interest (white line) are shown in (E). Asterisks indicate haustorial bodies. White bars correspond to 20µm. Pictures represent z-stacks of 15 confocal sections of each 2 µm increment.

## Discussion

ICR/RIP proteins are considered scaffold proteins in ROP signaling. Next to RICs, ICRs/RIPs might be key factors in branching of ROP signaling in plants. It appears that so far most described downstream functions of ROPs are mediated through either RIC or ICR/RIP proteins. ICRs/RIPs contain a characteristic QWRKAA motif in the CC2 domain, which was previously described as the motif responsible for ROP interaction (Lavy et al., 2007). Our results support this, since only full length RIPb and truncations containing this motif interacted with RACB and were subcellularly recruited by CA RACB (Fig. 3, Fig. 4, Supplemental Fig. S5, S6). The CC2 domain is part of all predicted ICRs/RIPs form *A. thaliana, O. sativa, B. distachyion* and *H. vulgare*. All identified ICRs/RIPs from these four species also contain a conserved QD/EEL motif located in an N-terminal CC1 domain (Supplemental Fig. S1). The function of this motif, however, remains more elusive. Although the CC1 domain is important for microtubule localization of RIPb (Fig. 4), amino acid exchanges in the QDEL motif did not result in loss of microtubule association (Supplemental Fig. S6).

Phylogenetic analyses show that both rice and Brachypodium possess putative orthologs of each of the three barley ICRs/RIPs, implying possible conserved function of the ICRs/RIPs in grasses (Supplemental Figs. S1 and S2). However, the five ICR/RIP proteins of Arabidopsis show no clear phylogenetic relation to the grass ICRs/RIPs. It would be interesting to see, whether Arabidopsis and grass ICRs/RIPs have similar functions, or may have evolved in different directions as the little sequence conservation suggests. In this context, it is noteworthy, that barley RIPa has functions reminiscent of Arabidopsis RIP3/ICR5/MIDD1 in microtubule organization, as recently shown (Hoefle et al., 2020). Hence, although these proteins are little conserved when compared between mono- and dicots, they may share at least some conserved functions in ROP signaling.

For this study, we focused on a possible RACB signaling mechanism via ICR/RIP proteins during the interaction of barley and *Bgh*. Barley RIPb interacts with CA and wild type RACB in yeast, supporting that it is a potential downstream interactor of RACB. Over-expression of RIPb but not of RIPa or RIPc increased penetration rate of *Bgh* into transformed epidermal barley cells (Fig. 1, Supplemental Fig. S4A). RIPb silencing had no significant effect on the interaction between epidermal cells and *Bgh* (Fig. 1B). This might be due to residual transcript or protein amounts of RIPb after transient knockdown or due to convergence in RACB downstream signaling which could compensate for the lack of RIPb during the interaction. For instance RIC171 might act as an alternative downstream interactor of RACB (Schultheiss et al., 2008), and it is possible that even more interactors of RACB are involved, because ROP proteins are considered signaling hubs (Nibau et al., 2006). Hence silencing of only one signaling branch might not have a significant effect on the interaction, whereas over-expression could support a certain RACB downstream branch and therefore has an effect. Additionally, RACB is not the only barley ROP that can support fungal penetration success (Schultheiss et al., 2003), and hence even RACB-independent ROP signaling could compensate for RACB-RIPb functions in RIPb-silenced cells.

RIPb shows diverse subcellular localizations. Next to cytosolic localization, we observed localization at the plasma membrane and at the microtubule cytoskeleton (Fig. 2). The N-terminal CC1 domain is necessary but not sufficient for microtubule localization, since the RIPbVaCC2 truncation lacking the CC1-domain did not localize to microtubules. The CC1 domain alone did also not show microtubule localization. The central Va domain alone was insufficient for microtubule association but it appeared to be required for both microtubule association and for RIPb-RIPb interaction (Fig. 4). BiFC experiments further suggested that the RIPb-RIPb interaction takes mainly place at microtubules (Fig. 5). Interestingly, truncated versions of RIPb, which contain the Va domain, did not induce susceptibility when over-expressed, whereas RIPbCC1 and particularly RIPbCC2 induced susceptibility, similar to or much stronger than the full length protein. We therefore speculate that dimerization or oligomerization of RIPb at microtubules might have a regulatory purpose, potentially by sequestration of inactive RIPb. However, interpretation of these results is complicated because the amount of expressed protein truncations also differed and could partially explain the differences in efficacy. Nevertheless, over-expression of the RIPbCC2 domain resulted in a very strong increase in susceptibility of barley epidermal cells to *Bgh*. Lavy et al. (2007) showed that the QWRKAA motif in the CC2 domain of Arabidopsis AtICR1/AtRIP1 is not only necessary for ROP interaction, but also for the interaction with the downstream interactor AtSEC3, indicating that the CC2 domain might be able to fulfill the signaling function of AtICR1/AtRIP1. In a follow-up publication by the same group they also show that the last 40 amino acids of AtRIP1 are also required for interaction with CMI1 (Ca™-dependent modulator of ICR1) (Hazak et al., 2019). This might be comparable to RIPbCC2, if RIPbCC2 would bind in addition to RACB a yet to be identified downstream protein. By contrast, over-expression of the CC2 domain of HvRIPa did not result in a significant increase in susceptibility (Supplemental Fig. S4B), and therefore this effect appears specific for RIPb. RIPbCC2 was able to interact with RACB in yeast and *in planta* (Fig. 4, Supplemental Fig. S7). Furthermore, RIPb did not localize to the cell periphery anymore without the CC2 domain (RIPbCC1Va) even in presence of CA RACB (Supplemental Fig. S6). This together suggests, that the CC2 domain of RIPb is responsible both for ROP interaction and for a function, which my take place at the plasma membrane.

The N-terminal CC1 domain of RIPb is required for microtubule association but might interact with signaling components as well. This could explain the susceptibility effect of over-expression of RIPbCC1, although the CC1 domain itself does not interact with RACB (Fig. 1C, Fig. 4C). Interestingly, the CC1 domain of Arabidopsis AtRIP3/ICR5/MIDD1 is required for interaction with KINESIN13A (Mucha et al., 2010). It could hence be that RIPb fulfills a dual function via different domains of the protein.

BiFC experiments showed interaction between RACB and RIPb at the microtubules and at the plasma membrane. Since RACB alone does not localize to microtubules (Schultheiss et al., 2003) it seems that RIPb is able to recruit RACB to microtubules when over-expressed. The interaction between the susceptibility-inducing CC2 domain and RACB on the other hand takes place at the plasma membrane (Supplemental Fig. S5, S6). These results suggest that RACB also recruits RIPb to the plasma membrane during susceptibility signaling and that recruitment of RACB to microtubules perhaps limits this effect. We speculate that in this experimental setup, recruitment of RACB to microtubules brings RACB into proximity of microtubule-located MAGAP1, which presumably inactivates RACB (Hoefle et al., 2011). This might explain why full length RIPb has a less strong effect on susceptibility when compared to RIPbCC2, which cannot recruit RACB to the microtubules. We found that protein levels of YFP-RIPbCC2 are higher than the levels of full length YFP-RIPb when transiently expressed in epidermal cells (Supplemental Fig. 6). Since both constructs are driven by a CaMV35S promotor, a different post-transcriptional regulation or protein turnover might be the most plausible explanation for this. The difference in protein levels can influence the effect that both proteins have, which would also confirm our notion, that RIPbCC2 might be a less regulated functional version of RIPb. However, since RIPbVaCC2 showed similar protein levels to RIPb, but had no influence on the outcome of the interaction between barley and *Bgh* it is unlikely that protein levels alone explain different efficacies of over-expression constructs.

We observed co-localization of RIPb and RACB and of RIPbCC2 and RACB at the site of fungal attack. In interactions where the fungus was able to penetrate the host cell, a ring of RIPb and RACB or CA RACB around the haustorial neck at the plasma membrane, was observed. However, we also observed signals at repelled penetration attempts around the formed papilla, indicating that accumulation of these two proteins alone is not sufficient to render all cells susceptible. RACB possesses a C-terminal CSIL motif, which is predicted to mediate protein prenylation at the cysteine residue, and is necessary for plasma membrane association and function in susceptibility (Schultheiss et al., 2003). Additionally, RACB has a polybasic stretch close to the C-terminus (Schultheiss et al., 2003) shown for other ROPs to be involved in lipid interaction (Platre et al., 2019) and a conserved cysteine at position C158, which is S-acylated in activated Arabidopsis AtROP6 (Sorek et al., 2017). Hence, lipid modification and interaction with negatively charged phospholipids together may bring activated RACB to specific membrane domains, to which it then recruits proteins that execute ROP signaling function. Phosphatidylserine and phophoinositides are often involved in defining areas of cell polarization in membranes for example during root hair and pollen tube tip growth (Helling et al., 2006; Kusano et al., 2008; Platre et al., 2019) and ROPs are known to moderate the phosphorylation pattern of phosphoinositides during polarization (Kost et al., 1999). We hence speculate that localization of ROP signaling components at the site of interaction reflects domains of enriched negatively charged phospholipids.

The exact effect of RACB-RIPb signaling on the interaction remains unknown so far. However, the finding that Arabidopsis ICRs/RIPs interact with proteins of the exocyst complex and KINESIN13A opens the possibility that barley ICRs/RIPs also modify the cytoskeleton or membrane trafficking, both being key to resistance and susceptibility in powdery mildew interactions (Hückelhoven and Panstruga, 2011; Dörmann et al., 2014), although at least for RIPb we found no strong evidence that the microtubule cytoskeleton might be affected (Supplemental Fig S9). Together, our data support a hypothesis according to which RIPb is localized at microtubules from which it is recruited to RACB signaling hotspots at the plasma membrane by activated RACB. There it might interact with further proteins of the RACB signaling pathway but also with RACB-independent factors to facilitate fungal entry into barley epidermal cells. The fact that the putative fungal effector ROPIP1 binds RACB and destabilizes barley microtubules (Nottensteiner et al., 2018) adds another level of complexity, on which ROPIP1 may foster release of RIPb from microtubules for its function in susceptibility.

## Conclusions

Over the last years, the impact of susceptibility factors for plant – pathogen interactions has become more and more obvious. Barley susceptibility factor RACB might be a key player in cellular polarization during fungal invasion. Here we identified RIPb as a potential downstream interactor of activated RACB in susceptibility. RACB and RIPb might be involved in fine-tuning of cell polarization in advantage of the fungus. It will be important to identify further interactors of RIPb and of its strongly susceptibility-supporting CC2 domain. This may establish a deep understanding of the components and mechanisms of subcellular reorganizations in the cell cortex, which support the biotrophic parasite *Bgh* in accommodation of its haustorium in an intact epidermal cell.

## Material and Methods

### Biological Material

Barley (*Hordeum vulgare*) cultivar Golden Promise was used in all experiments. Plants were grown under long day conditions with 16h of light and 8h in the dark with a relative humidity of 65% and light intensity of 150 µM s^™1^ m^™2^at a temperature of 18°C.

Powdery mildew fungus *Blumeria graminis* f.sp. *hordei* race A6 was cultivated on wild type Golden Promise plants under the conditions described above and inoculated by blowing spores into a plastic tent that was positioned over healthy plants or transformed leaf segments.

### Cloning procedures

*HvRIPb* (*HORVU1Hr1G012460*) was amplified from cDNA using primers Ripb-EcoRI_fwd and Ripb-BamHI_rev (Supplemental Tab. 1) introducing EcoRI and BamHI restriction sites, respectively. *HvRIPa* (HORVU3Hr1G087430) was amplified from cDNA using primers RipaXbaI_fwd and RipaXbaI_rev introducing XbaI restriction sites at 5’ and 3’ ends. *HvRIPc* (HORVU3Hr1G072880) was amplified from cDNA using primers RipcXbaI_fwd and RipcPstI_rev introducing restriction sites for XbaI at the 5’ end and for SalI at the 3’ end. The amplified products were ligated into the pGEM-T easy vector (Promega, Madison, WI, USA) by blunt end cloning according to the manufacturer’s instructions and sequenced. *HvRIPb* truncations spanning the following amino acids. *HvRIPbCC1* from amino acid 1 to 132, *HvRIPbVa* from amino acid 133 to 420 and *HvRIPbCC2* from amino acid 420 to 612. *HvRIPb* truncations for Yeast-Two-Hybrid were amplified from pGEM-T easy containing full length *RIPb* using primers with EcoRI and BamHI restriction sites. *RIPbCC1* was amplified using primers Ripb-EcoRI_fwd and RipbCC1BamHI_rev, *RIPbCC1Va* with primers Ripb-EcoRI_fwd and RipbVaBamHI_rev, *RIPbVa* with primers RipbVaEcoRI_fwd and RipbVaBamHI_rev, *RIPbVaCC2* with primers RipbVaEcoRI_fwd and Ripb-BamHI_rev and *RIPbCC2* with primers RipbC2EcoRI_fwd. Each reverse primer introduced a stop codon. For Yeast-Two-Hybrid assays *HvRIPb* and *HvRIPb* truncations were subcloned from the pGEM-T easy vector into pGADT7 and pGBKT7 plasmids (Clontech Laboratories) using the EcoRI and BamHI restriction sites. For over-expression constructs and constructs for protein localization the pUC18-based vector pGY1, containing a CaMV35S promotor was used. (Schweizer et al., 1999). From the pGEM-T easy vector, *HvRIPb* was further amplified with primers Ripb-XbaI_fwd and Ripb-SalI_rev, containing XbaI and SalI restriction site, respectively. Using those restriction sites *HvRIPb* was then ligated into the pGY1 plasmid and pGY1-YFP plasmid for N-terminal YFP fusion. *HvRIPa* and *HvRIPc* were subcloned form pGEM-T easy into pGY1 using the XbaI restriction site for *HvRIPa* and the XbaI and PstI restriction sites for *HvRIPc*. Over-expression construct for *HvRIPaCC2* was produced by introducing attB-attachment sites for Gateway cloning. For this, a first PCR was performed with primers GW1-RipaCC2_fwd and GW1-Ripa_rev using pGEM-T easy construct as template. A subsequent second PCR was performed using primers Gate2_F and Gate2_R to introduce attB attachment sites for Gateway cloning. The construct was then cloned by BP-clonase reaction using the Gateway BP Clonase™II (Invitrogen) into the pDONR223 entry vector (Invitrogen). From there *HvRIPaCC2* was cloned by LR-clonase reaction with Gateway LR Clonase™II (Invitrogen) into pGY1-GW, a modified pGY1 vector containing the gateway cassette. The pGY1-GW plasmid was constructed using the Gateway™Vector Conversion System (Invitrogen) according to the manufacturer’s instructions.

For BiFC, *HvRIPb* was amplified from the pGEM-T easy vector using the primer Ripb-SpeI_fwd and Ripb-SalI_rev with restriction sites for SpeI and SalI, respectively. The construct was then digested with SpeI and SalI and ligated into pUC-SPYNE(R)173 and pUC-SPYCE(MR) plasmid (Waadt et al., 2008) using these restriction sites.

A 538bp long RNAi sequence for *HvRIPb* was amplified, using primers RipbRNAi_fwd and RipbRNAi_rev, and introduced into the pIPKTA38 vector by blunt-end cloning using the SmaI restriction site (Douchkov et al., 2005). This plasmid was used as entry vector to clone the RNAi Sequence into the pIPKTA30N vector for double-strand RNA formation via Gateway LR Clonase ™ II (Invitrogen) reaction according to the manufacturer’s instruction.

All *HvRIPb* truncations were introduced into the pGY1-YFP plasmid for N-terminal YFP fusion using the following primer. For *HvRIPbCC1* primer Ripb-XbaI_fwd and RipbC1-SalI_rev, for *HvRIPbCC1Va* primer Ripb-XbaI_fwd and RipbVa-SalI_rev, for *HvRIPbVa* primer RipbVa-XbaI_fwd and RipbVa-SalI_rev, for *HvRIPbVaCC2* primer RipbVa-XbaI_fwd and Ripb-SalI_rev and for *HvRIPbCC2* primer RipbC2-XbaI_fwd and Ripb-SalI_rev. All forward primers introduce a XbaI restriction site and all reverse primer contain a SalI restriction site, which were used for the ligation into pGY1-YFP. The same products and restriction sites were used for ligation into the pGY1 vector except for *HvRIPbCC1Va*. For *HvRIPbCC1Va* primer GW-Ripb_fwd and GW1-RipbC1Va_rev was used for amplification followed by a second PCR with primers Gate2_F and Gate2_R to introduce attB attachment sites for Gateway cloning. The construct was then cloned by BP-clonase reaction using the Gateway BP Clonase™II (Invitrogen) into the pDONR223 entry vector (Invitrogen). From there *HvRIPbCC1Va* was cloned by LR-clonase reaction with Gateway LR Clonase™II (Invitrogen) into pGY1-GW.

### Transient transformation of barley cells

Barley epidermal cells were transiently transformed by biolistic particle bombardment using the PDS-1000/HE (Biorad, Hercules, CA; USA). For this 7d old primary leaves of barley were cut and placed on 0.8% water-agar. Per shot 302.5µg of 1µm gold particles (Biorad, Hercules, CA, USA) were coated with 1µg plasmid per shot. 0.5µg plasmid per shot was used for cytosolic transformation markers. After addition of plasmids to the gold particles, CaCl_2_ was added to a final concentration of 0.5M. Finally, 3µl of 2mg/ml Protamine (Sigma) were added to the mixture per shot. After incubation for half an hour at room temperature, gold particles were washed twice with 500µl ethanol. In the first step with 70% ethanol and in the second step with 100% ethanol. After washing, the gold particles were re-suspended in 6µl of 100% ethanol per shot and placed on the macro carrier for bombardment.

### Alignments and Phylogenetic Analysis

Sequences of Arabidopsis ICR/RIP proteins were used to identify barley ICRs/RIPs using the IPK Barley BLAST Server (https://webblast.ipk-gatersleben.de/barley_ibsc/viroblast.php). ICRs/RIPs from *Oryza sativa spp. Japonica* were identified using the BLAST tool on the Rice Genome Annotation Project (http://rice.plantbiology.msu.edu/home_faq.shtml (Kawahara et al., 2013)). ICRs/RIPs from *Brachypodium distachyon* were identified by BLAST search on EnsemblPLANTs (https://plants.ensembl.org/index.html). The Alignment of ICR/RIP protein sequences was done with ClustalO (https://www.ebi.ac.uk/Tools/msa/clustalo/) and displayed with Jalview (jalview 2.10.5). A phylogenetic maximum likelihood tree was generated, using the PhyML tool in the program seaview (v4.7).

### Determination of Susceptibility

Transiently transformed barley leaves were inoculated with *Bgh* 24 h after bombardment for over-expression constructs and 48 h after bombardment for gene silencing constructs. 24 h after inoculation penetration rate into the transformed cells was determined by fluorescence microscopy as described before (Hückelhoven et al., 2003).

### Protein localization and Protein – Protein Interaction *in planta*

Localization of HvRIPb and co-localization of HvRIPb and HvRACB were determined by transiently transforming barley epidermal cells with plasmids encoding fluorophore fusion proteins. Imaging was done with a Leica TCS SP5 microscope equipped with hybrid HyD detectors. CFP was excitated at 458nm and detected between 465nm and 500nm. YFP was excitated at 514nm and detected between 525nm and 570nm. Excitation of mCherry and RFP was done at 561nm and detection between 570nm and 610nm.

For ratiometric quantification of BiFC experiments Mean Fluorescence Intensity (MFI) was measured over a region of interest at the cell periphery. Background signal was subtracted and ratio between YFP and mCherry signal was calculated. At least 25 cells were analyzed per construct for each experiment. Images were taken 24 hours to 48 hours after transformation by particle bombardment.

To evaluate the microtubule to cytosol signal ratio, ten cells per construct were measured. MFI in each cell was measured either on cytosolic strands or along microtubules on three different regions of interest each, in single imaging plains. Average MFI was calculated for cytosol and microtubules, respectively. Afterwards ratio between average microtubule signal and average cytosolic signal was calculated.

### Yeast Two-Hybrid assays

For targeted yeast two-hybrid assays, *HvRIPb* and its truncations were introduced into pGADT7. Introduction of *HvRACB* into pGBKT7 was described in Schultheiss et al. (2008). Constructs were transformed into yeast strain AH109 following the small-scale LiAc yeast transformation procedure from the Yeast Protocol Handbook (Clontech, Mountain View, CA, USA).

### RNA extraction and semiquantitative PCR (sq-PCR)

RNA was extracted from barley tissue using the TRIzolTM-Reagent by Invitrogen according to the manufacturer’s instructions. 1µg of RNA was reverse transcribed with the QuantiTect Reverse Transcription Kit (Qiagen, Hilden, Germany) according to the manufacturer’s instructions.

For semiquantitative PCR, 2µl of cDNA transcribed from RNA of pealed epidermis from barley leaves, were used. Samples were taken from leaves 24h after inoculation with *Bgh*, or from uninoculated leaves of the same age. A 209bp fragment of *RIPa* was amplified with an annealing temperature (T_a)_ of 58°C with primers Ripa_sqPCR4_fwd and Ripa_sqPCR5_rev (Supplemental Tab1). For *RIPb* a 181bp fragment was amplified with a T_a_ of 56°C using primers Ripb_sqPCR9_fwd and RIPb_sqPCR10_rev. For *RIPc* a 168bp fragment was amplified at T_a_ 58°C using primers Ripc_sqPCR4_fwd and Ripc_sqPCR5_rev. As control *HvUbc* was amplified at T_a_ 61°C using primers HvUBC2_fwd and HvUBC2_rev.

## Supporting information

Merged Supplements

## Acknowledgements

The authors would like to thank Johanna Hofer for her technical assistance and Dr. Caroline Höfle, Dr. Michaela Stegmann and Clara Igisch for their support of this work (all TU Munich, Phytopathology).

