## Supplementary material for "ROP INTERACTIVE PARTNER b interacts with RACB and supports fungal penetration into barley epidermal cells": Merged Supplements

### SUPPLEMENTAL MATERIAL

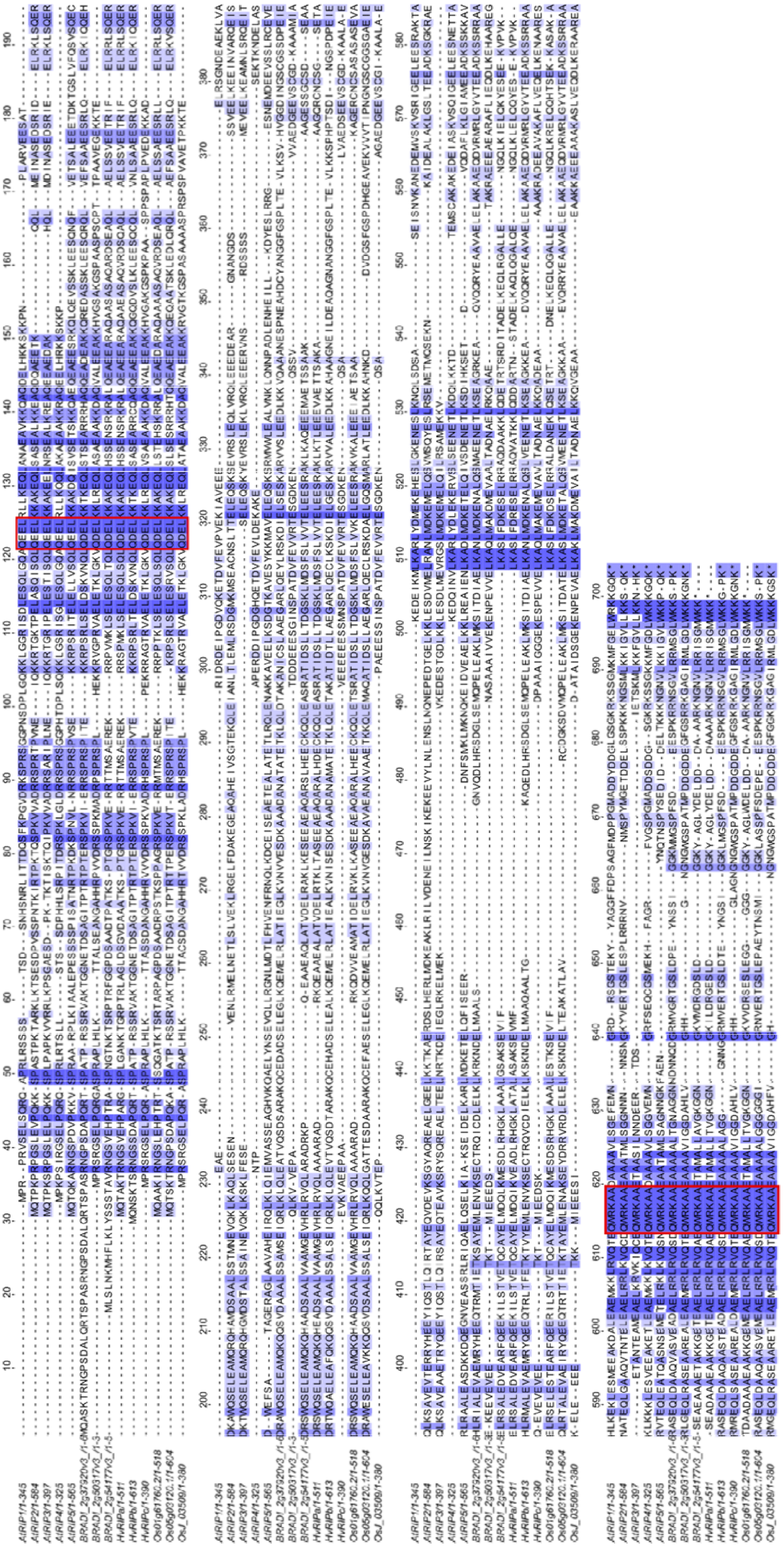

**Supplemental Fig. S1.** (A) Alignment of amino acid sequences of barley ICR/RIP proteins with ICR/RIP proteins from *Arabidopsis thaliana* and rice. Alignment was carried out with ClustalO and displayed with jalview (jalview 2.10.5). Color intensity relates to sequence identity. Red rectangles mark the QD/EEL and the QWRKAA motif. AtRIP1=AtICR1, AtRIP2=AtICR2, AtRIP3=AtMIDD1=AtICR5; AtRIP4=AtICR4, AtRIP5=AtICR3.

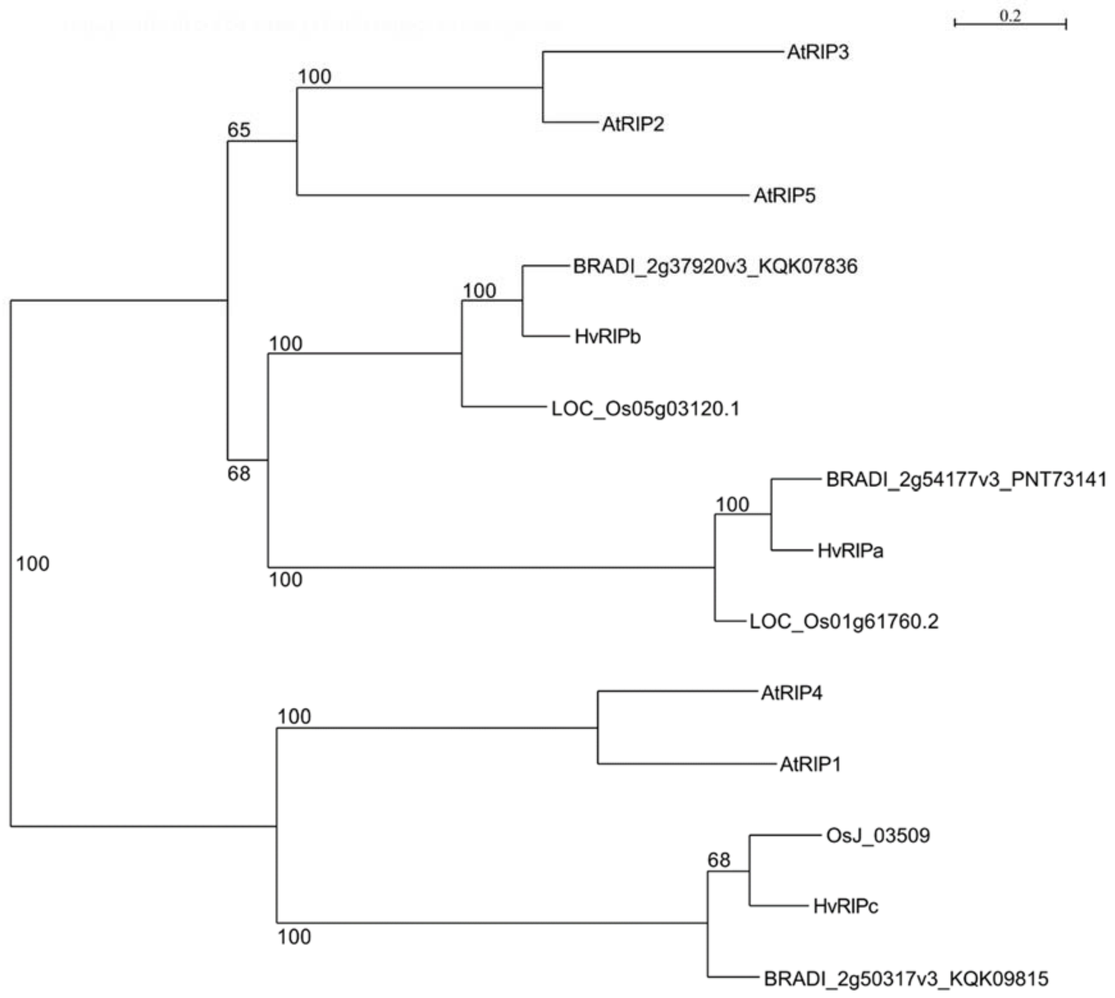


**Supplemental Fig. S2.** Phylogenetic relationship of ICR/RIP proteins. A phylogenetic maximum likelihood tree was generated, including three additional ICR/RIP proteins from *Brachypodium distachyon* using the PhyML tool in the program seaview (v4.7). AtRIP1=AtICR1, AtRIP2=AtICR2, AtRIP3=AtMIDD1=AtICR5; AtRIP4=AtICR4, AtRIP5=AtICR3.

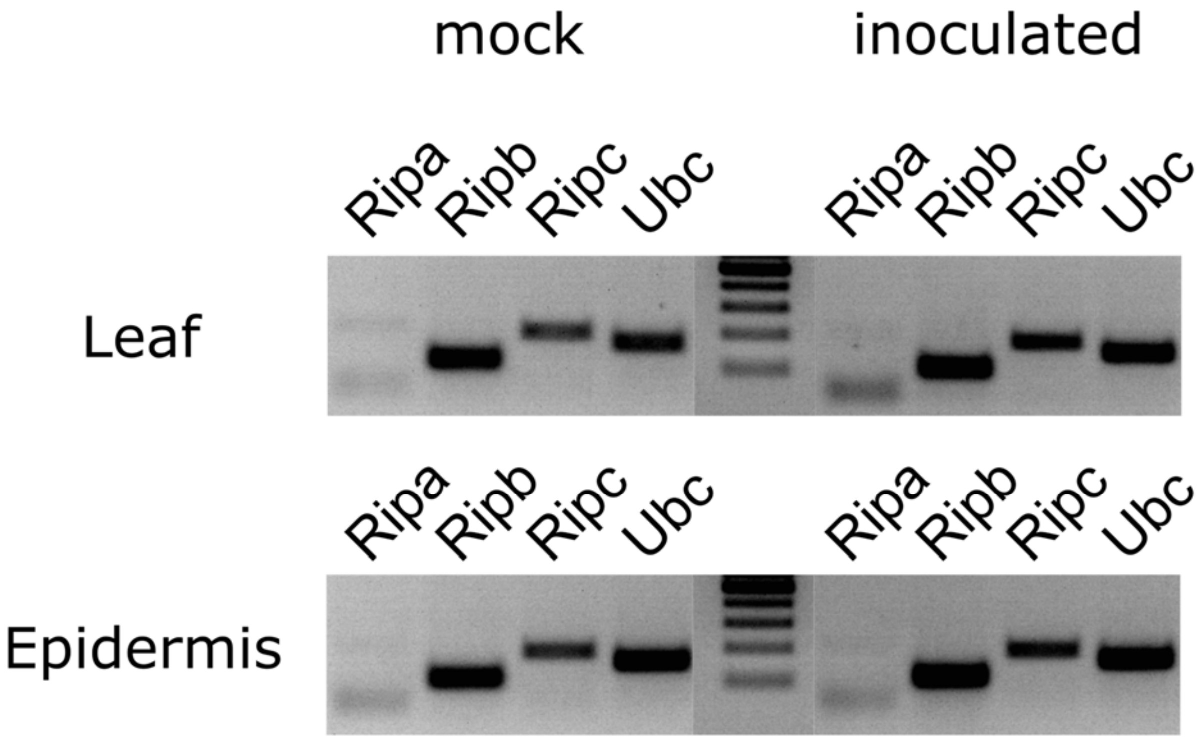


**Supplemental Fig. S3**. Semiquantitative PCR shows transcription levels of *HvRIPa*, *HvRIPb* and *HvRIPc*. Samples were taken from whole barley leaves or leaf epidermal layers, either inoculated with *Bgh* or not. As control, the constitutively expressed housekeeping *HvUbc* gene was used. Equal amounts of cDNA were used to perform sqPCR. For each *ICR/RIP* gene, primers were designed amplifying around 200 bp from parts close to the 5' seqeunce. Expected amplicon sizes are 209bp for *RIPa*, 109bp for *RIPb*, 188bp for *RIPc*, and 156bp for *Ubc*.

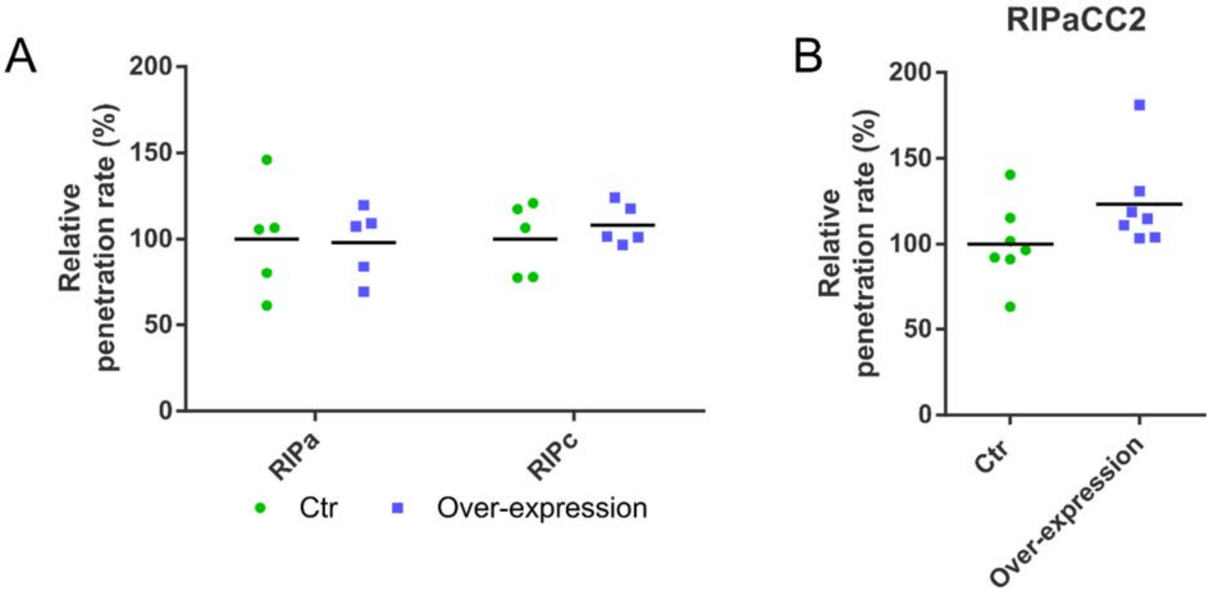


**Supplemental Fig. S4**. Effect of RIPa and RIPc on the interaction of barley and *Bgh* was tested by biolistic transformation of epidermal cells of 7 days old barley plants. The penetration rate of *Bgh* into the transformed cells was determined 24 h after inoculation. Over-expression constructs for *RIPa* and *RIPc* (A) as well as an over-expression construct of *RIPaCC2* (B) were introduced. As control, the respective expression empty vectors were used. Values represent the mean values of results of individual experiments (n≥5) relative to the mean of the respective control set as 100 %.

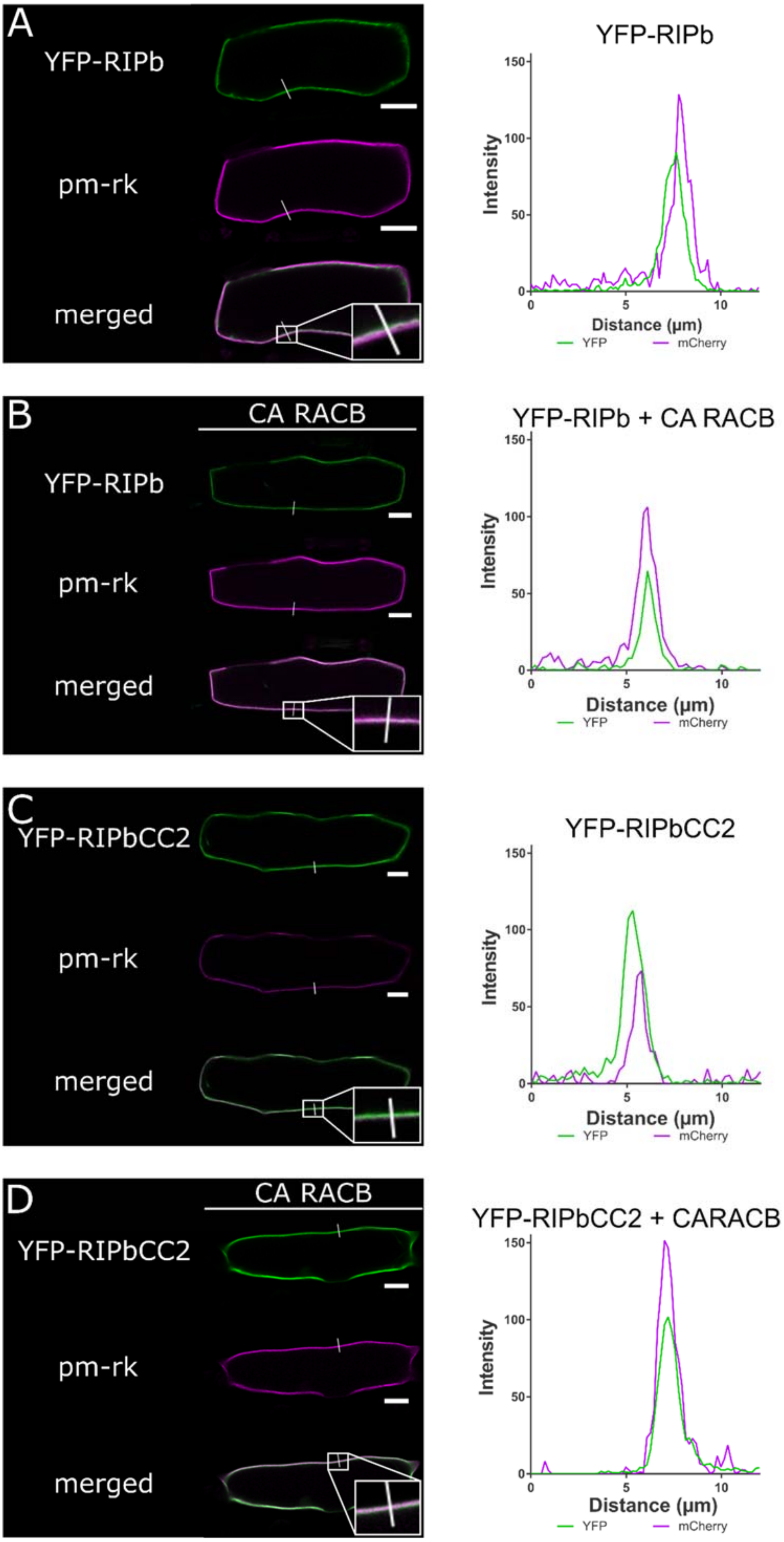
A
YFP-RIPb
pm-rk
merged
YFP-RIPb
Intensity
Distance (µm)
YFP
mCherry
B
CA RACB
YFP-RIPb
pm-rk
merged
YFP-RIPb + CA RACB
Intensity
Distance (µm)
YFP
mCherry
C
YFP-RIPbCC2
pm-rk
merged
YFP-RIPbCC2
Intensity
Distance (µm)
YFP
mCherry
D
CA RACB
YFP-RIPbCC2
pm-rk
merged
YFP-RIPbCC2 + CARACB
Intensity
Distance (µm)
YFP
mCherry

**Supplemental Fig. S5.** RIPb gets recruited to the plasma membrane by RACB. YFP-RIPb (A, B) and YFP-RIPbCC2 (C, D) were transiently expressed in barley epidermal cell together with the plasma membrane marker pm-rk (Nelson et al., 2007; Weis et al., 2013). Additionally, YFP-RIPb and YFP-RIPbCC2 were co-expressed with CA RACB (B, D). Signal profile was measured over a linear ROI (white line) and displayed in the respective graph on the right hand side. White bars correspond to 20µm.

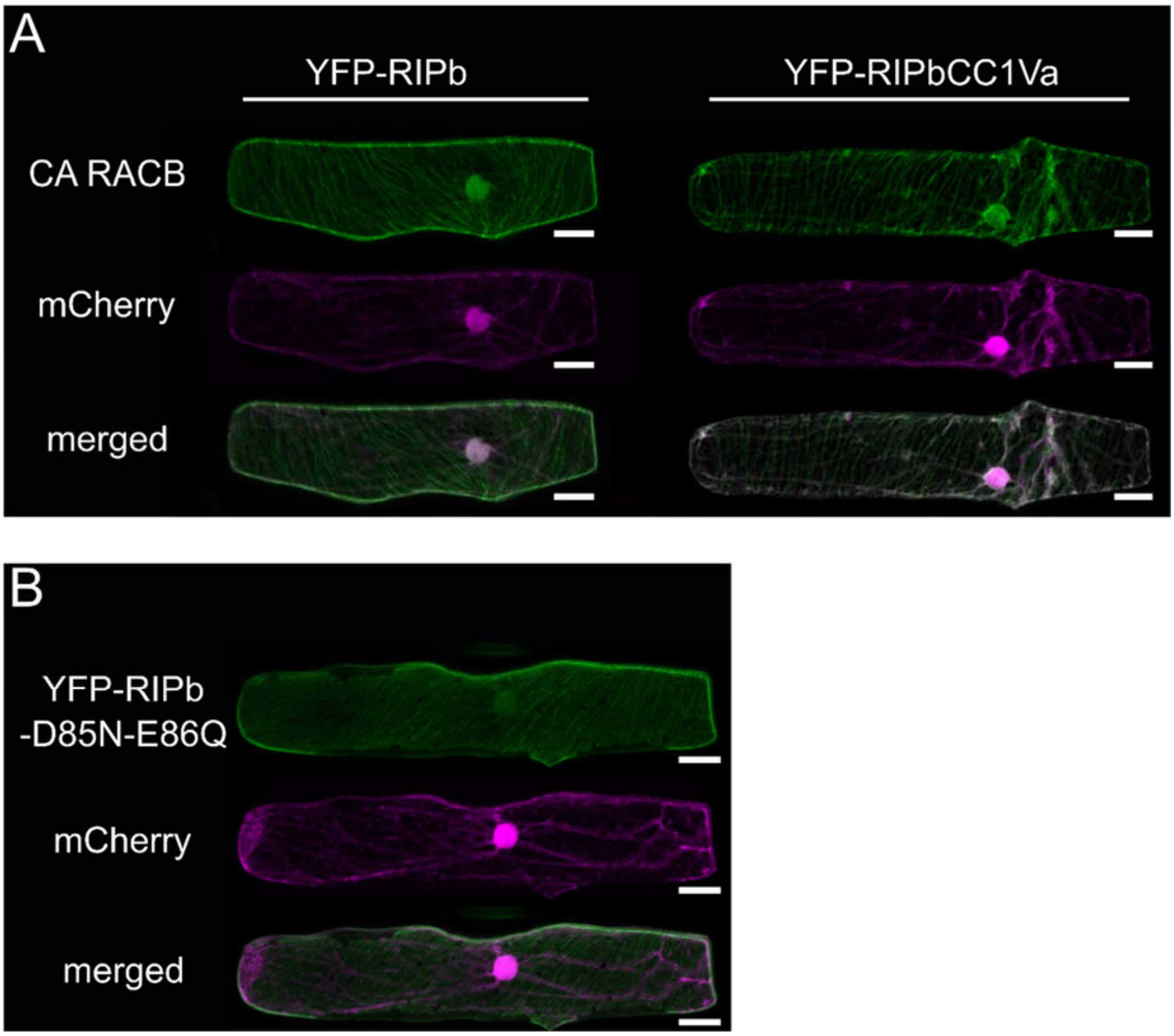


**Supplemental Fig. S6**. RIPbCC1Va cannot be recruited to the cell periphery by RACB. (A) Single epidermal cells were transiently transformed by particle bombardment with CA RACB, mCherry and either YFP-RIPb or YFP-RIPbC1Va. (B) Mutation D85N and E86Q were introduced into RIPb and a YFP-fusion protein was transiently expressed in single epidermal cells. White bars correspond to 20µm.

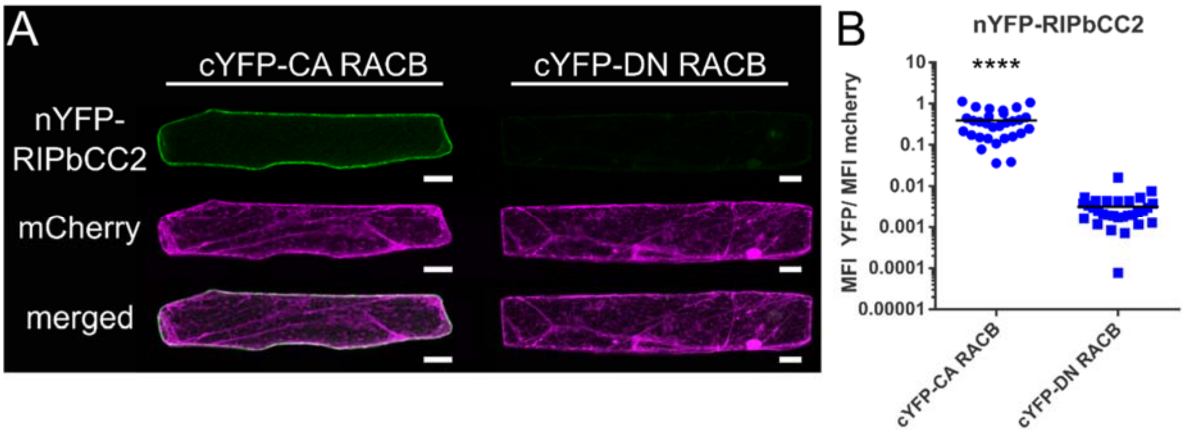


**Supplemental Fig. S7**. The CC2 domain of RIPb interacts with RACB *in planta*. Single epidermal cells of barley leaves were transiently transformed by particle bombardment with fusion proteins expressing split-YFP constructs for BiFC. (A) nYFP-RIPbCC2 was either co-expressed with cYFP-CA RACB or cYFP-DN RACB. mCherry served as transformation control. White bars correspond to 20µm. (B) For quantification of YFP complementation images were taken with constant settings and signal intensity (Mean Fluorescence Intensity, MFI) was measured over a region of interest at the cell periphery. The ratio between YFP and mCherry signal was calculated (n=30).

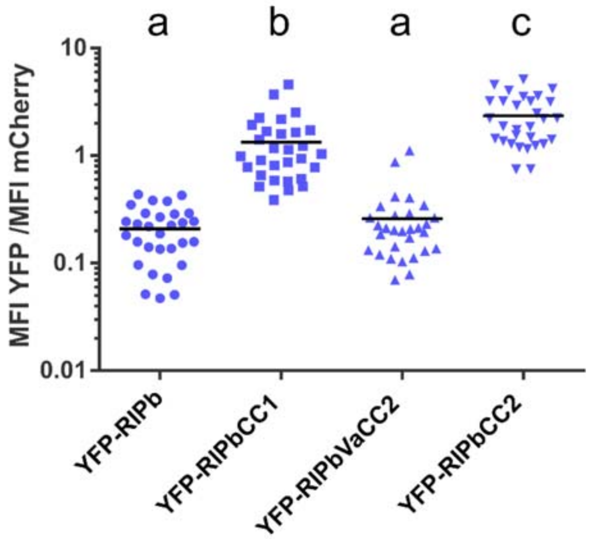


**Supplemental Fig. S8.** Stability of RIPb. YFP fusion proteins were transiently expressed in single barley epidermal cells together with mCherry as an internal reference in order to measure expression levels. Transformed cells were imaged by confocal microscopy and Mean Fluorescence Intensity was measured for the upper half of the cell and calculated relative to the internal mCherry reference. For each construct at least 30 cells were measured. Letters indicated significance by one-way ANOVA (Tukey's multiple comparison test).

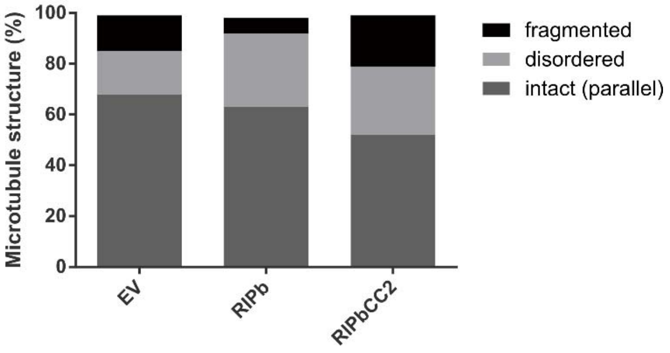


**Supplemental Fig. S9.** Cortical microtubule organization in barley epidermal cells was assessed 2d after transiently co-expressing microtubule marker RFP-MAGAP1-Cterm), cytosolic transformation marker GFP and indicated constructs. Graph shows percentage of barley epidermal cells displaying particular microtubule order categories indicated on the right. EV = empty overexpression plasmid. Total number of cells for each sample > 140 in 3 independent reps.

*Supplemental Table 1* Primer list

| Name | Sequence (5'-3') | Restriction sites /attachment sites | Product |
|---|---|---|---|
| **Ripb-EcoRI_fwd** | AGAATTCATGCAGAACTCAAAAACCAGTAG | EcoRI | *RIPb*<br>*RIPbCC1* |
| **Ripb-BamHI_rev** | TGGATCCGGTCTCATGAGCTTCTTCAC | BamHI | *RIPb*<br>*RIPbCC2* |
| **Ripb-XbaI_fwd** | TCTAGATATGCCGAGATCCAG | XbaI | *RIPb*<br>*RIPbCC1* |
| **Ripb-SalI_rev** | AGTCGACCGGTCTCATGAGCT | SalI | *RIPb*<br>*RIPbCC2* |
| **Ripb-SpeI_fwd** | TACTAGTTTCATGCAGAACTCAAAAACCAGTAG | SpeI | *RIPb* |
| **RipbRNAi_fwd** | ATCTAGACAGAGGCACGAAGGTGCCAAGCAC | XbaI | *RIPb-RNAi* |
| **RipbRNAi_rev** | TGTCGACCTTCAGGCATTCTTGCAACCGGGC | SalI | *RIPb-RNAi* |
| **RipbC1-SalI_rev** | TGTCGACTCAGAGATTGACAAGCTGGCA | SalI | *RIPbCC1* |
| **RipbVa-XbaI_fwd** | ATCTAGATATGTCAGCAGCAGAGGAGTCC | XbaI | *RIPbVa*<br>*RIPbVaCC2* |
| **RipbVa-SalI_rev** | TGTCGACTCATTCGCTCAGCCCGTCTG | SalI | *RIPbVa*<br>*RIPbCC1Va* |
| **RipbC2-XbaI_fwd** | ATCTAGACGAAATGCAGCCGGAGC | XbaI | *RIPbCC2* |
| **GW-Ripb_fwd** | GCAGGCTCAGGAATGCAGAACTCAAAAACCAGTAG | | *RIPbCC1Va* |
| **GW1-RipbC1Va-st_rev** | GAAAGCTGGGTCCTCATTCGCTCAGCCCGTCTG | | *RIPbCC1Va* |
| **Gate2_F** | GGGACAAGTTTGTACAAAAAAGCAGGCTCA | attB1 | |
| **Gate2_R** | GGGGACCACTTTGTACAAGAAAGCTGGGTC | attB2 | |
| **RipaXbaI_fwd** | TCTAGATATGCAGACAGCCAAGACAAG | XbaI | *RIPa* |
| **RipaXbaI_rev** | TCTAGATCATTTCTTCCACATTCCACTG | XbaI | *RIPa* |
| **RipcXbaI_fwd** | TCTAGATATGCAGAACTCAAAAACC | XbaI | *RIPc* |
| **RipcPstI_rev** | TCTGCAGTCACCTTCACTTGTTGCCC | PstI | *RIPc* |
| **RipbC1BamHI_rev** | TGGATCCTCAGAGATTGACAAGCTGGCAC | BamHI | *RIPbCC1* |
| **RipbVaEcoRI_fwd** | AGAATTCTCAGCAGCAGAGGAGTCC | EcoRI | *RIPbVa*<br>*RIPbVaCC2* |
| **RipbVaBamHI_rev** | TGGATCCTCATTCGCTCAGCCCGTCTG | BamHI | *RIPbCC1Va*<br>*RIPbVa* |
| **RipbC2EcoRI_fwd** | AGAATTCGAAATGCAGCCGGAGC | EcoRI | *RIPbCC2* |
| **GW1-RipaCC2_fwd** | GCAGGCTCAATGCAGGACGACGCGAGAACG | | *RIPaCC2* |
| **GW-Ripa_rev** | GAAAGCTGGGTCTCATCATTTCTTCCACATTCCACTG | | *RIPaCC2* |
| **Ripa_sqPCR4_fwd** | GCCAAGACAAGGAATGGCTC | | *RIPa* |
| **Ripa_sqPCR5_rev** | GAGAGCTTCATGGGTGACCT | | *RIPa* |
| **Ripb_sqPCR9_fwd** | CCCAGTTACTGAGAAGAAGCG | | *RIPb* |
| **Ripb_sqPCR10_rev** | CAGCTTCAACGACACATCCTG | | *RIPb* |
| **Ripc_sqPCR4_fwd** | GCTGCCAGAGAAGAGGCG | | *RIPc* |

| | | | |
|---|---|---|---|
| **Ripc_sqPCR5_rev** | TTGGCGCCGACATGCTTC | | *RIPc* |
| **HvUBC2_fwd** | TCTCGTCCCTGAGATTGCCCACAT | | *UBC* |
| **HvUBC2_rev** | TTTCTCGGGACAGCAACACAATCTTCT | | *UBC* |
